# Conformational Pathways of Translational T-box Riboswitch Governing tRNA Recognition for Gene Regulation

**DOI:** 10.1101/2025.07.25.666911

**Authors:** Dibyendu Mondal, Govardhan Reddy

## Abstract

The availability of various amino acids is essential for the synthesis of proteins during translation. In bacteria, T-box riboswitches play a critical role in regulating the concentrations of amino acids by sensing and decoding the aminoacylation status of tRNAs for precise control of amino acid biosynthesis and transport pathways. We used computer simulations to probe the mechanism of tRNA recognition by a translational T-box riboswitch using the *iles* T-box riboswitch as a model system. We show that the T-box aptamer populates three distinct thermodynamic states – undocked, predocked, and docked – depending on the relative orientation between its three stems (stem I, II, and IIA/B). The relative population of the three states is strongly dependent on the Mg^2+^ concentration. In the undocked and predocked states, the freely moving stem I catches the tRNA from the solution using the ‘fly-casting’ mechanism. Subsequently, the tRNA-bound stem I docks to stem II in a specific orientation with the help of noncanonical intra- and inter-backbone hydrogen bonds, which facilitate tRNA interaction with the discriminator domain to detect the aminoacylation state of tRNA and regulate the gene. Since T-box riboswitches are prime targets in designing antibiotics, our proposed mechanism of tRNA recognition by T-box reveals multiple critical sites in the riboswitch that can be targeted for developing new antibiotics. Interestingly, molecules acting as antibiotics exist targeting one of the proposed sites. These results also have broad implications for understanding the role of specific RNA-RNA interactions in various cellular processes.

## Introduction

Precise control of the concentrations of amino acids is essential for cellular functions, because amino acids are critical components in protein synthesis during translation and are also actively involved in numerous essential metabolic pathways.^1–5^ In bacteria, imbalances in amino acid levels disrupt cellular processes such as cell division, cell wall formation, cell growth and metabolism, and quorum sensing.^6^ Cells employ multiple mechanisms to regulate amino acid concentrations, including control of biosynthesis pathways via feedback inhibition of key enzymes, modulation of amino acid transporters, and metabolic degradation or recycling.^7–10^ Among these, RNA-based regulatory methods such as riboswitches have emerged as sophisticated strategies, where structured RNA elements sense amino acid availability to directly modulate gene expression.^11,12^ Riboswitches are RNA structures that bind small molecules, ions, or metabolites, triggering changes in gene expression to respond precisely to the cellular metabolic state.^13–18^

The T-box riboswitches recognize specific tRNA molecules to regulate genes that encode for aminoacyl-tRNA synthetases and proteins involved in amino acid biosynthesis and transport.^19,20^ The T-box riboswitches are grouped into four distinct classes (I-IV) based on variations in their subdomain architecture.^19,21,22^ Classes I and II typically regulate transcription termination, and classes III and IV mainly control translation initiation.^23^ Several high-resolution crystal structures have elucidated the architecture of different T-box variants.^24–27^ The class III translational *ileS* T-box riboswitch is a single-stranded noncoding RNA composed of a decoder domain and a discriminator domain. The decoder domain consists of stem I (SI), stem II (SII), and stem IIA/B (Ψ). The discriminator domain consists of stem III (SIII) and anti-sequestrator (AntiS) module (Fig. 1a, b, S1).^25,27^ T-box riboswitches are promising targets for the development of RNA-based antibacterial strategies because their sequences and modular structure are highly conserved in multiple bacterial species, and there are also no human homologs of these riboswitches.^28–33^

**Figure 1:**
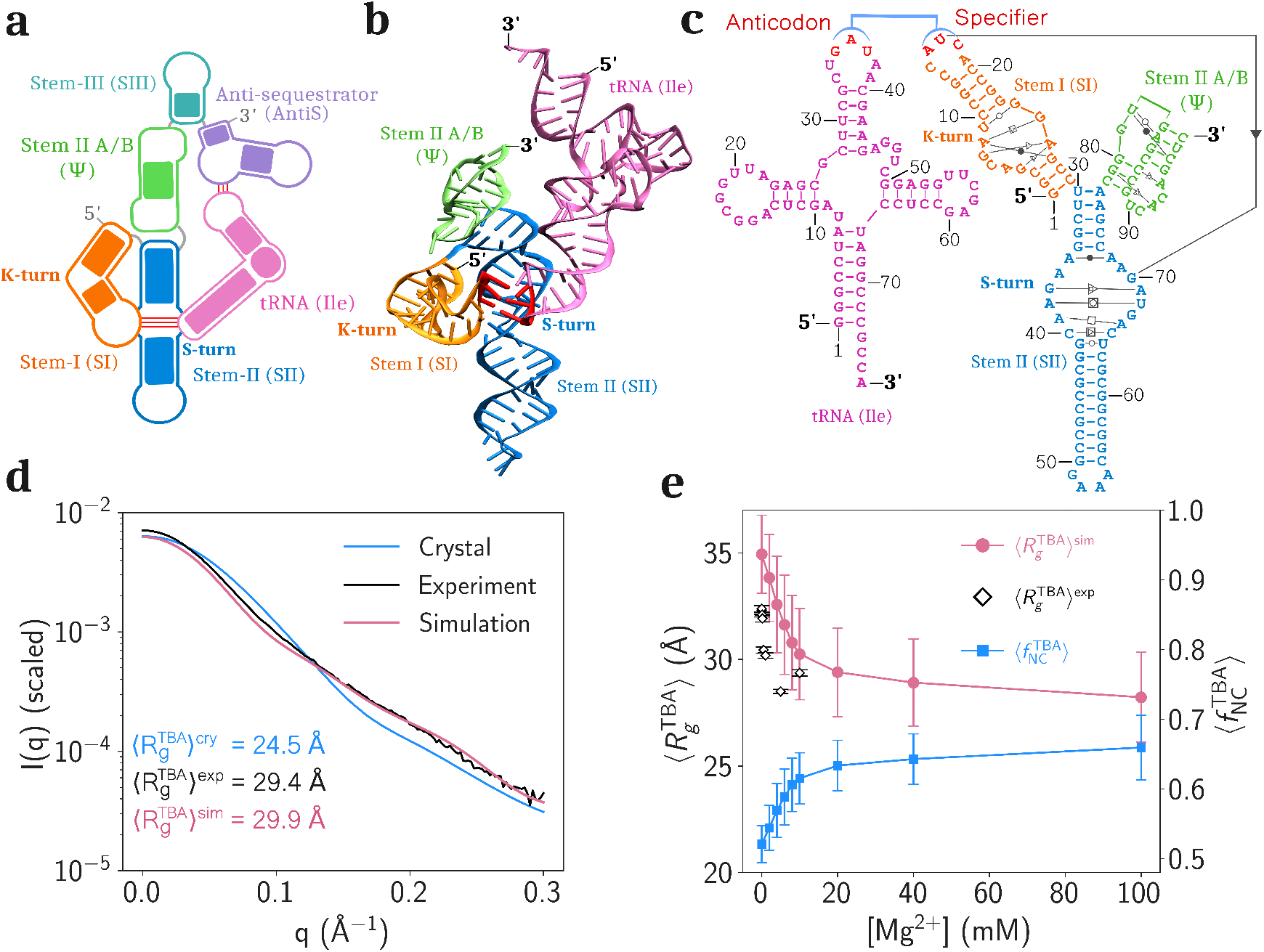
**(a)** Schematic of a T-box riboswitch bound to tRNA. The tRNA and the various subdomains of the riboswitch are shown. **(b)** Crystal structure of the decoder domain of the *ileS* T-box riboswitch aptamer (TBA) (PDB ID: 6UFM).^27^ Specifier and Anticodon sequences are highlighted in red. The stem III and Anti-sequestrator domains are absent in this crystal structure. **(c)** The secondary structure of TBA and tRNA. Noncanonical base pairs are in Tables S2, S3, S4, and for TBA they are denoted by Leontis-Westhof symbols.^46,47^ **(d)** SAXS profile of tRNA bound TBA. The experimental data (black) is from the work of Niu et al.^28^ The scattering intensities from the simulation trajectories (mauve) and the crystal structure (blue) are calculated using CRYSOL.^43^ **(e)** Average radius of gyration 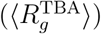 and average fraction of native contact 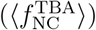 of TBA as a function of [Mg^2+^].

Crystal structures of *ileS* T-box riboswitch from different bacteria species show that in the native *holo* state, stem I with a K-turn kink docks onto stem II at the S-turn motif, forming the primary tRNA binding site (PDB IDs: 6UFM, 6UFH) (Fig. 1b, S1).^25,27^ The tRNA is recognized via base pairing between its Anticodon and the Specifier codon on stem I. The docked tRNA also interacts with the AntiS domain, which senses its aminoacylation status. Three-color single-molecule Förster resonance energy transfer (smFRET) experiments of the *ileS* T-box riboswitch demonstrated that the decoder domain adopts a heterogeneous ensemble of conformations involved in multistep tRNA recognition.^23^ smFRET studies on T-box mutants highlighted the roles of different subdomains and the influence of Mg^2+^ ions on tRNA binding.^28^ Recent smFRET experiments on full-length T-box revealed a two-step aminoacylation-dependent discrimination mechanism mediated by the discriminator domain.^34^ Collectively, the T-box riboswitch selectively binds its cognate tRNA, assesses its aminoacylation state, and triggers sequestrator or anti-sequestrator formation accordingly. Despite this progress, several fundamental questions remain: (i) What are the structures of the intermediate states populated during T-box folding? (ii) How does tRNA binding alter the folding landscape of T-box? (iii) What is the role of Mg^2+^ ions in the tRNA recognition and detection of the aminoacylation state of the tRNA by the T-Box riboswitch? (iv) What is the role of the S-turn in stem II on tRNA recognition and gene regulation?

Experimentally, it is challenging to simultaneously probe the T-box folding, tRNA binding, and the role of Mg^2+^ ions in these processes. It is extremely difficult to study short-lived transient intermediate states critical for tRNA recognition using experiments. Even in all-atom molecular dynamics (MD) simulations, it is challenging to sample the full conformational landscape because of the large size of RNA, the slow kinetics of folding and binding, and the complexity of the underlying free-energy surface. To overcome these limitations, we used the three-interaction-site (TIS) coarse-grained (CG) RNA model and molecular dynamics (MD) simulations to study the tRNA recognition by the decoder module of the *Nocardia farcinica ileS* T-box aptamer domain (TBA) (PDB ID: 6UFM).^27,35^

We validated the CG RNA model using the small-angle X-ray scattering (SAXS) and the smFRET experimental data on TBA.^23,28^ We show that TBA populates multiple states depending on the relative orientation between different arms in the T-box structure. The relative population of these states strongly depended on the Mg^2+^ concentration. We further show that TBA catches the freely diffusing tRNA in solution using the ‘fly-casting’ mechanism. The orientation of the tRNA docked to the TBA is critical to verifying its aminoacylation state, and noncanonical hydrogen bonds between TBA and tRNA play an important role in stabilizing the correct orientation of the docked tRNA. We finally discussed the implications of our results in targeting T-Box riboswitches for discovering new antibiotics.

## Methods

### Coarse-Grained MD Simulations

Several coarse-grained (CG) RNA models have been developed to address a variety of RNA-related questions.^36–41^ In this work, we used the three interaction site (TIS) RNA model^35,36^ to investigate the folding behavior of the *ileS T-box* riboswitch.^27^ All CG simulations were carried out using the OpenMM molecular simulation toolkit.^42^ Details on the TIS model and simulation protocols are provided in the Supporting Information.

### Calculation of Scattering Profile from Simulation

The scattering profile from the simulation was calculated using the CRYSOL package.^43^ We used the simulation data of *apo* TBA at [Mg^2+^] = 10 mM and 298 K. We converted the CG structures to energy-minimized all-atom constructs using the TIS2AA package^44^ and supplied the structures to CRYSOL for the calculation of the theoretical scattering intensities.

### Fraction of Native Contacts

The fraction of native contacts (NC) between a particular set of nucleotides (*λ* = TBA, RBD) in a conformation *i*, is computed using 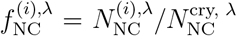 where 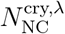 and 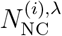 are the number of native contacts present between the nucleotides belonging to set *λ* in the crystal structure and the *i*^th^ conformation, respectively.^45^ See Data Analysis in Supporting Information for further details.

### Local Ion Concentration Around Backbone

The local concentration of an ion M^*n*+^, around a specific backbone phosphate bead is given by,^35,45^ 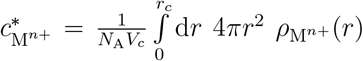, where 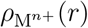 is the number density of ion M^*n*+^ at a distance *r* from the backbone phosphate bead under consideration, *V*_*c*_ is the spherical volume with a cutoff radius *r*_*c*_ (= 5 Å), and *N*_A_ is the Avogadro number. See Data Analysis in Supporting Information for further details.

## Results

### Validation of Mg^2+^-induced Folding of T-box Aptamer Domain

Physiologically abundant cations, such as Mg^2+^, play a critical role in TBA folding and its binding to tRNA since both their backbones are negatively charged due to the phosphate groups.^28^ We performed MD simulations using the TIS RNA model^36^ to study tRNA binding to TBA in Mg^2+^ concentration ([Mg^2+^]) ranging from 2 to 100 mM at a temperature *T* = 298 K. In these simulations, a constant background concentration of K^+^ ([K^+^]) was maintained at 150 mM (physiological concentration).

To validate the TIS model, we compared the experimental SAXS scattering profiles of *apo* TBA with those computed from simulations. Both sets of profiles were obtained under similar thermodynamic conditions ([Mg^2+^] = 10 mM, *T* = 298 K) and in the absence of tRNA^28^ (see Methods). The scattering profile computed using the crystal structure is in poor agreement with the experimental profile (Fig. 1d).^28^ Also the average radius of gyration calculated from the experimental scattering data 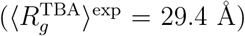 is also greater than the value obtained using the crystal structure 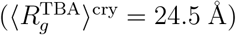 suggesting that there are structural differences between *apo* TBA and native tRNA-bound TBA conformations.^28^ The scattering profile computed from the simulations is in better agreement with the experiments, suggesting that the ensemble of conformations obtained from the simulations of the *apo* TBA is representative of the experimental ensemble and is different from the crystal structure. Moreover, the radius of gyration computed from the simulations 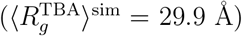 is in excellent agreement with the value measured in experiments 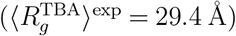. The larger 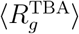 value of *apo* TBA compared to the crystal structure shows that its conformations are swollen without tRNA binding.

The average radius of gyration, 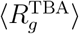, as a function of [Mg^2+^] shows that 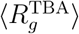 decreased sharply from 35 Å to 29 Å as [Mg^2+^] increased from 0 to 20 mM, indicating rapid compaction of TBA from the unfolded state to the folded state (Fig. 1e). However, even at 100 mM, the 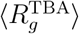 remained greater than that of 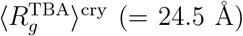, suggesting that even at high [Mg^2+^], the *holo*-TBA structure (crystal-like structure) is not stable without tRNA binding. The 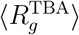 from the simulations is in reasonable quantitative agreement with the 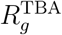 measured in the experiments (Fig. 1e).^28^

To confirm that the crystal-like structure is not stable without tRNA binding, we com-puted the average fraction of native contacts 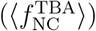 as a function of [Mg^2+^]. We observed that 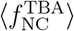 increased from 0.52 to only 0.65 with an increase in [Mg^2+^] from 0 to 20 mM, and reached a plateau for [Mg^2+^] *>* 20 mM (Fig. 1e). This suggests that increasing Mg^2+^ shifts the TBA population toward a native-like state, but it does not dominantly populate the native state without tRNA binding.

### TBA Populates Three States in the Absence of tRNA Binding

To understand the role of Mg^2+^ in the folding of TBA, we mapped the FES onto 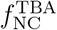 for different [Mg^2+^] (Fig. 2a, S2) (see Data Analysis in SI). At a low [Mg^2+^] (2-4 mM), an undocked state (U1) is populated at 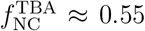. At higher [Mg^2+^] (6–100 mM), two additional states are populated − the state 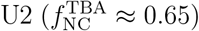 and the state 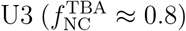.

**Figure 2:**
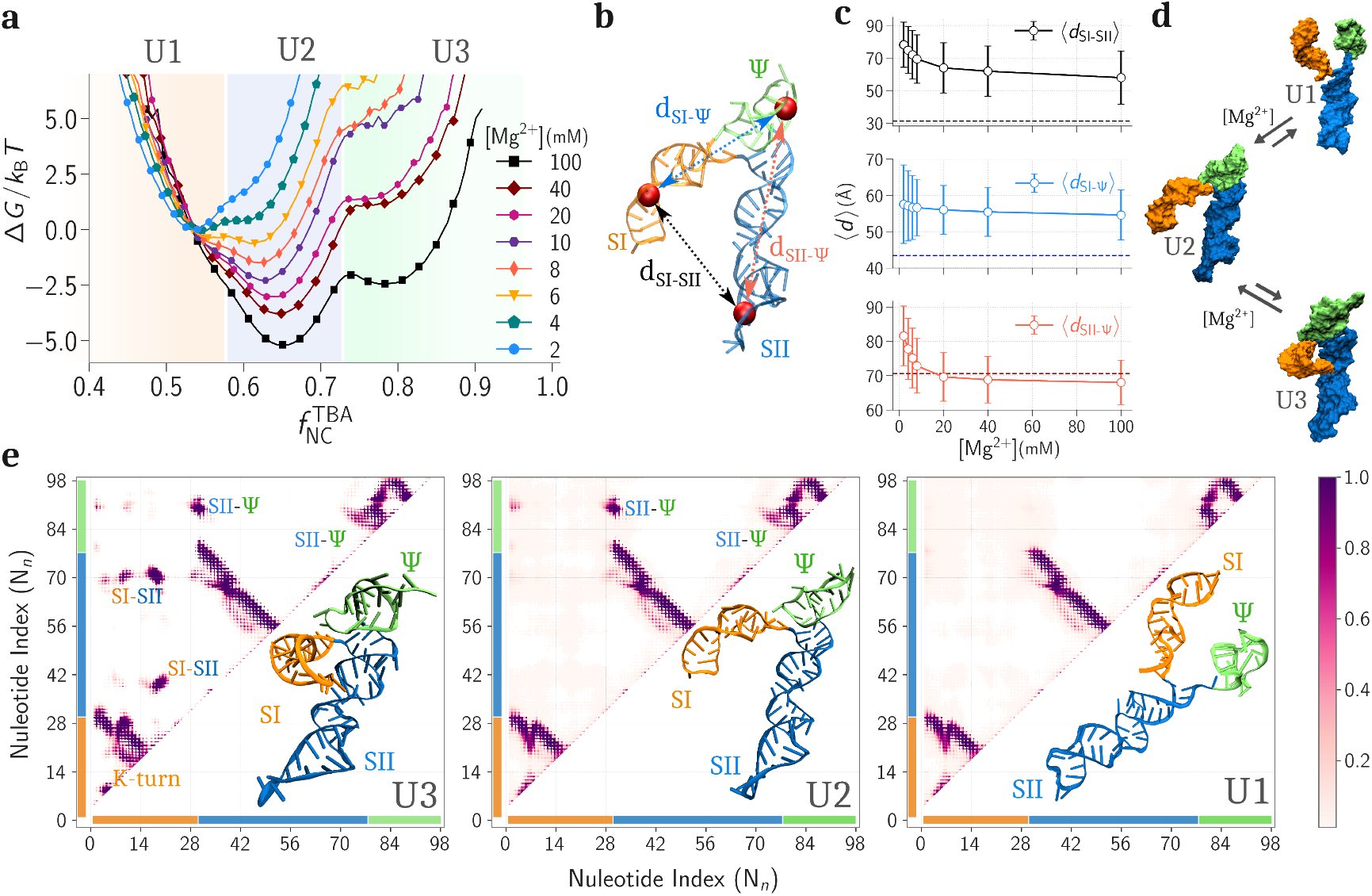
**(a)** Free energy surface of TBA projected on the average fraction of native contacts 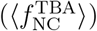 for different [Mg^2+^] at 298 K. Three states U1, U2, and U3 are highlighted in orange, blue and green shaded regions, respectively. **(b)** The distances between stems are calculated using the distances between the center of mass of nucleotides G12, G13, C20, C21 for stem I, C49, G50, C55, G56 for stem II, and A84, G85, C96, U97 for Ψ (shown in red beads). **(c)** The average distances (*d*_SI-SII_, *d*_SI-Ψ_, and *d*_SII-Ψ_) between stems as a function of [Mg^2+^]. The corresponding distances in the crystal structure are shown by the dashed lines. **(d)** The equilibrium between three states of TBA. **(e)** The average contact maps for states U3, U2, and U1 showing the probabilities for different tertiary contact formation. The representative structures are shown in the inset. The color bars on the axes are a guide for the nucleotide sequences for different stems.

To quantify structural transitions, we measured the distances between three stems (Fig. 2b). The average distance between stems I and II, ⟨*d*_SI-SII_⟩, decreased from 78 Å to 58 Å as [Mg^2+^] increased from 2 to 100 mM, indicating structural compaction (Fig. 2c,S3). But, even at high [Mg^2+^] (= 100 mM), the ⟨*d*_SI-SII_⟩ (= 58 Å) is still quite large compared to that of the crystal structure (≈ 31 Å), implying that without tRNA binding, the TBA preferably remains in the undocked states (U1 and U2). The average distance between stems I and Ψ, ⟨*d*_SI-Ψ_⟩, shows a slight decrease from 58 Å to 55 Å but deviates from the distance in the crystal (≈ 43 Å), suggesting the absence of the docked state. Interestingly, the average distance between stems II and Ψ, ⟨*d*_SII-Ψ_⟩, decreased from 81 Å to 68 Å with increasing [Mg^2+^] and is close to the crystal distance (≈ 71 Å) implying Mg^2+^-induced stabilization of native-like coaxial stacking of stems II and Ψ. The equilibrium between these states was confirmed by the multiple transitions between them (Fig. 2d, S2). smFRET studies identified three thermodynamic states in the TBA folding process by analyzing the distances between stem I and stem II. These experiments show that at low [Mg^2+^], TBA folds to an intermediate state but cannot form the native docked state, which is consistent with our results.^23,28^

For analyzing the structural differences between the three states, we computed the average contact map for each state (U1, U2, and U3). The contact map of U3 shows multiple inter-stem tertiary contacts, indicating a native-like docked state. These contacts are mostly dominated by the intra-backbone (sugar-sugar, base-sugar) hydrogen bonds between stems I and II known as the ribose-zipper interactions (Fig. S4).^23,27^ The contact map of U2 shows that the SI-SII contacts responsible for the docking of stems I and II are absent. However, the SII-Ψ contacts remain intact, pointing to stable coaxial stacking between stem II and Ψ. We refer to U2 as a predocked state. In U1, both SI-SII and SII-Ψ contacts are absent, suggesting a completely open undocked conformation. In the absence of tRNA, the FES shows that TBA populates three states (U1, U2, and U3), and their relative population depends on [Mg^2+^].

### tRNA Binding to the TBA Exhibits Kinetic Partitioning

To investigate the FES for tRNA binding to TBA, we performed simulations of TBA with a tRNA molecule in the simulation box for [Mg^2+^] = 4, 8, 12, 16, and 20 mM with a fixed background [K^+^] = 150 mM. For each [Mg^2+^], we performed 8 independent simulations. First, we discuss the data for [Mg^2+^] = 20 mM. In the presence of tRNA, 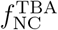 as a function of time shows that TBA samples all three states − undocked (U1), predocked (U2), and docked (U3) (Fig. S5a (cyan)). To verify tRNA binding to TBA, we computed the fraction of native contacts between tRNA and TBA in the tRNA binding domain (RBD) of TBA,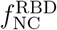 (Table 1 and Fig. S6). The low 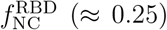 indicates unbound-undocked states, the high 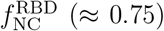 indicates bound-docked states, and the values in between indicate intermediate states (Fig. 3a).

**Table 1:** Description of nucleotides of sub-parts of tRNA and TBA

| Region | Nucleotides |
| --- | --- |
| Anticodon | G35, A36, and U37 |
| Specifier | A16, U17, and C18 |
| tRNA-binding-domain (RBD) | TBA: A16-A19, A36-A39, A69-A72, and tRNA: G35-A38 |

**Figure 3:**
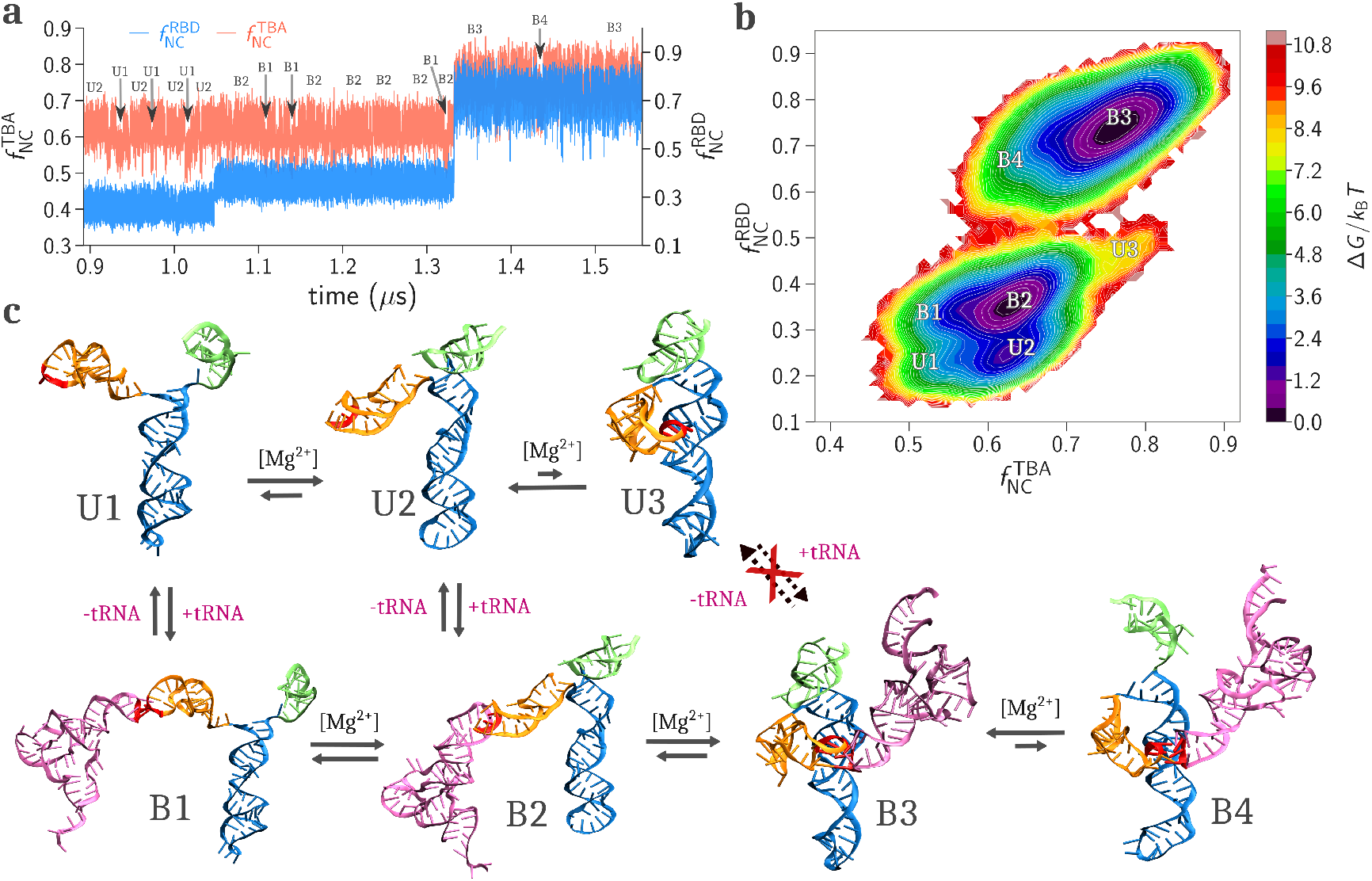
**(a)** A representative trajectory showing the fraction of native contacts of the RNA binding domain 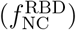 and the fraction of native contact of TBA 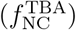 as a function of time at [Mg^2+^] = 20 mM. The transitions between different states are highlighted with arrows. All other possible transitions are shown in Fig. S5. **(b)** The FES projected onto 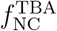 and 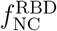 shows seven distinct states for [Mg^2+^] = 20 mM. **(c)** The kinetic network with representative structures from all the seven basins. The relative sizes of the forward and backward arrows indicate whether the equilibrium is left-shifted or right-shifted.

The 2D FES projected on 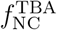 and 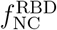 shows seven distinct states − three tRNA unbound states (U1, U2, and U3) and four tRNA bound states (B1, B2, B3, and B4) (Fig. 3b). All the representative structures for each state are shown in Fig. 3c, and all the representative transitions observed between the states are shown in Fig. S5. States B1 (undocked-bound) and B2 (predocked-bound) are the partially bound states where the tRNA binds to U1 and U2, respectively, through the Specifier loop of stem I (Fig. 3c and 1a). The tRNA-bound stem I remains undocked in states B1 and B2. B3 (docked-bound) is the native state where stem I is docked to the S-turn region of stem II and tRNA is bound to the RBD. In state B4 (uncapped-docked-bound), stems I and II are docked to the tRNA, but Ψ is not coaxially stacked to stem II.

### tRNA Binding with TBA Depends on [Mg^2+^]

To investigate the effect of [Mg^2+^] on tRNA binding to TBA, we performed simulations of TBA in the presence of tRNA at different [Mg^2+^] (Fig. 4, S7). At low [Mg^2+^] (≈ 4 mM), only four states (U1, U2, B1, and B2) are populated, where U1 and B1 were the dominant states. This suggests that primary binding of the tRNA Anticodon to the Specifier sequence of stem I is possible even at low [Mg^2+^]. Upon increasing [Mg^2+^] to 8 mM, the four states, U1, U2, B1, and B2, are populated, but no stem I-stem II docking (U3) event was observed. At [Mg^2+^] = 12 mM, the equilibrium population of U2 and B2 increased, indicating that Ψ is coaxially stacked with stem II. At this point, the other three states (U3, B3, and B4) also begin to populate, suggesting that the docking of the tRNA-bound stem I to the S-trun of stem II requires a critical [Mg^2+^] ≈ 12 mM. At [Mg^2+^] = 20 mM, B3 becomes more stable. For [Mg^2+^] *>* 20 mM, B3 is highly stable and is in equilibrium with a very low population of B4. The formation of U3 is less frequent at high [Mg^2+^] due to the strong Anticodon-Specifier binding affinity, leading to a dominant U2 to B2 transition compared to the U2 to U3 transition. We did not observe any transition from U3 to B3 because of the low population of U3 and the lower accessibility of the Specifier sequence as it is docked at the S-turn in U3.

**Figure 4:**
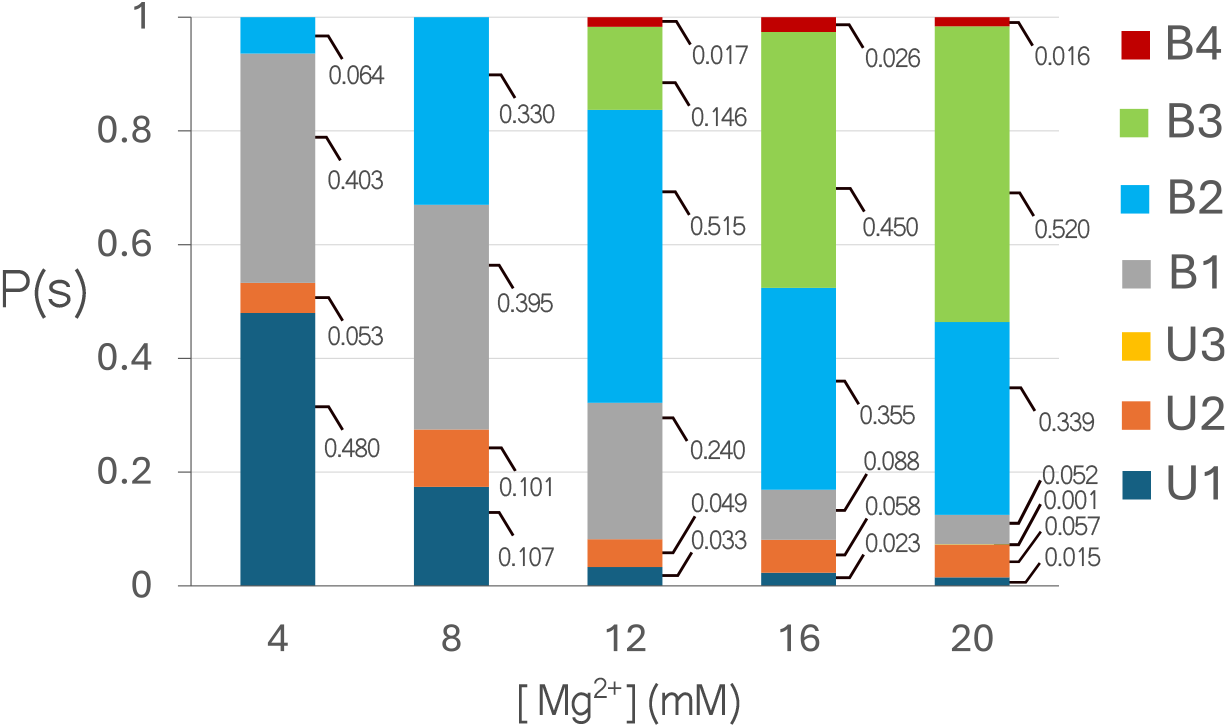
The equilibrium probabilities of states U1, U2, U3, B1, B2, B3, and B4 as function of [Mg^2+^].

### Mg^2+^ Stabilizes the Tertiary Structures Critical for tRNA Docking

The equilibrium between TBA states depends on [Mg^2+^] (Fig. 2a, 4). tRNA binding to TBA is through direct interaction between nucleotides and also through indirect interactions between nucleotides mediated by Mg^2+^. We computed the local Mg^2+^ concentration, 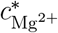 around each of the negatively charged backbone phosphate beads of the tRNA-unbound states of TBA (U1, U2, and U3) (see Methods).^35^ In the absence of tRNA, each state exhibits a distinct 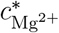 profile (Fig. 5a). U1 does not show significant peaks, while the states U2 and U3 show strong Mg^2+^ condensation around the nucleotides C28-U30 (junction of stem I and II) and C87-C90 (Ψ region), which mediates the tertiary contacts involved in the coaxial stacking between Ψ and stem II (Fig. 5a,b). The docked state, U3, shows additional peaks around the Specifier nucleotides (A16-C20) on stem I and S-turn nucleotides (A69-A71, A38-C40) on stem II. This indicates that the docking of stem I to stem II is coupled to Mg^2+^ binding. A strong Mg^2+^ peak is also observed around the nucleotides A8-C10 within the K-turn of stem I, indicating that Mg^2+^-induced bending of stem I is important to stabilize the docked conformation, which aligns with the previous experiments.^28,48^

**Figure 5:**
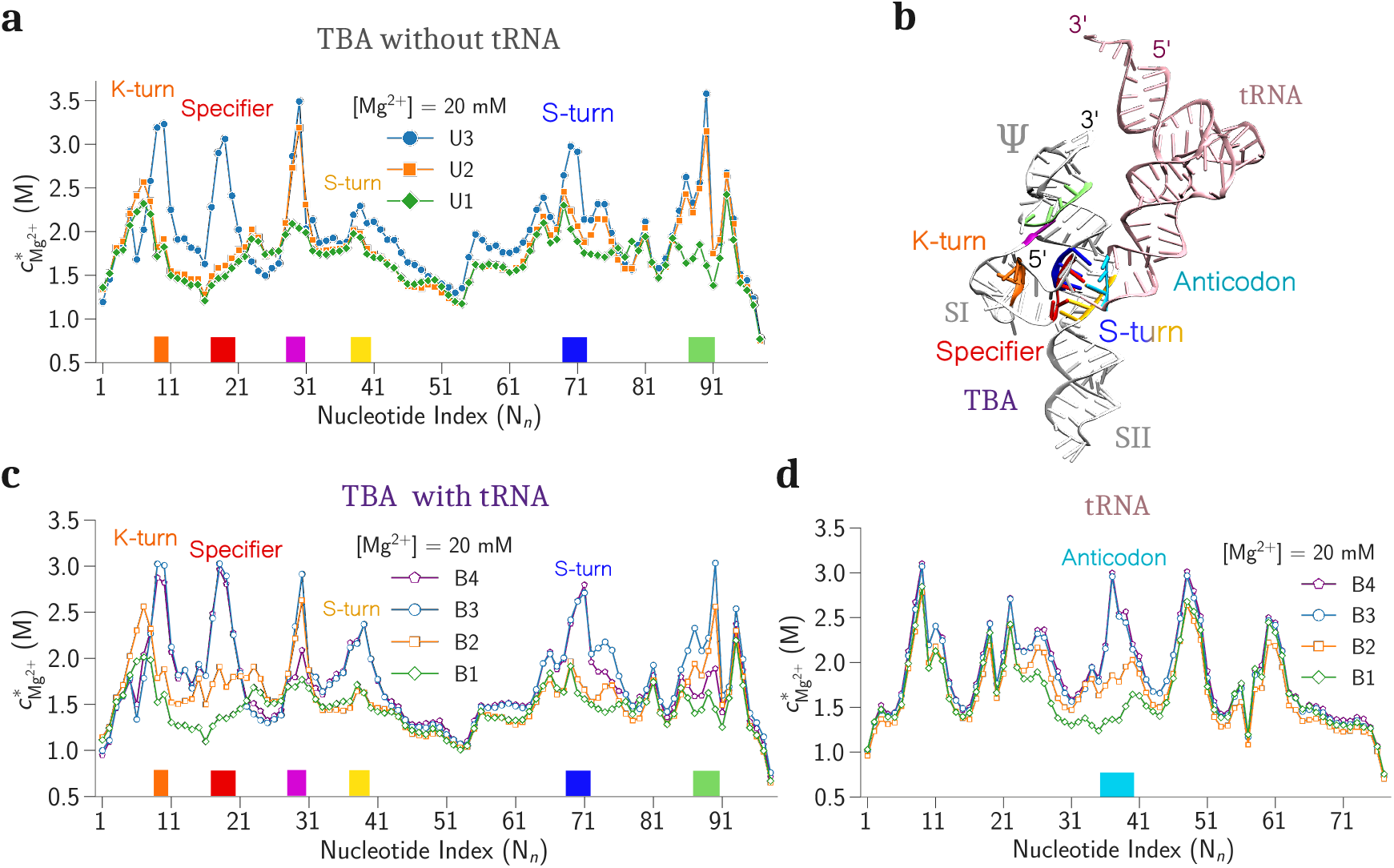
**(a)** Local concentrations of Mg^2+^ ions, 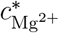 around the phosphate beads of TBA for states U1, U2, and U3. **(b)** Locations of the nucleotides with significant Mg^2+^ condensation on the crystal structure. **(c)** The 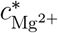 around the phosphate beads of TBA for states B1-B4. **(d)** The 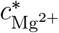 around the phosphate beads of tRNA for states B1-B4.

To study the role of Mg^2+^ in tRNA binding, we computed 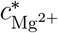 for all the tRNA-bound states (B1, B2, B3 and B4) (Fig. 5c). The 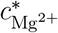 profiles of B1, B2, and B3 closely match U1, U2, and U3, respectively, with some differences. In B3, nucleotides G37-C40 and A69-A71 (S-turn motif) show increased Mg^2+^ condensation compared to U3, suggesting a Mg^2+^-induced stabilization of tRNA binding to RBD. Furthermore, compared to B1, B2 exhibits more Mg^2+^ condensation near the K-turn region and the Specifier-Anticodon binding region, which stabilizes the primary tRNA binding to TBA (Fig. 5b,c). The 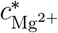 profile of B4 resembles B3 but lacks peaks at C28-U30 and C87–C90, indicating disruption of Mg^2+^-induced coaxial stacking between Ψ and stem II. The 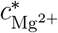 profiles for tRNA show increasing condensation only in the Anticodon region from B1 to B4. This suggests that tRNA binding to stem I is weakly Mg^2+^-dependent, while the docking of the tRNA-bound stem I to the S-turn of stem II is strongly dependent on Mg^2+^.

### Inter-backbone H-bonds between tRNA and S-turn are Critical in Stabilizing Docked states

Simulations showed that in the absence of tRNA, the stem I-stem II docked state (U3) was moderately stable even at very high [Mg^2+^] (≈ 100 mM). However, when tRNA is bound to stem I, the stability of the bound-docked state (B3) increases significantly, and it is the most stable state even at [Mg^2+^] = 20 mM. The crystal structure (PDB ID: 6UFM) showed that stem I docks to the S-turn via the sugar-sugar hydrogen bonds. In addition to the Specifier-Anticodon Watson-Crick base pairs, the tRNA also forms multiple inter-backbone hydrogen bonds (base-sugar, sugar-sugar, phosphate-sugar hydrogen bonds) with the S-turn (Table S4 and Fig. S8).^27^ These inter-backbone hydrogen bonds are almost invariant to sequence alteration. Therefore, to probe the importance of these hydrogen bonds, we did not allow these hydrogen bonds to form in the simulation by switching off their interaction potential. We observed that without the tRNA−S-turn inter-backbone hydrogen bonds, the stability of the tRNA-bound docked state (B3) decreased significantly even at high [Mg^2+^] (≈ 100 mM) (Fig. S9). Thus, the tRNA−S-turn inter-backbone H-bonds play a pivotal role, along with Mg^2+^ ions, in stabilizing the native state (B3).

We emphasize that in the presence of the tRNA−S-turn inter-backbone H-bonds, we never observed a state where the tRNA directly binds to the S-turn motif without initially binding to the Specifier in stem I. This indicates that although the tRNA−S-turn inter-backbone hydrogen bonds are pivotal in stabilizing the native bound-docked states (B3, B4), this stabilization is only possible if tRNA binding to the S-turn occurs with the assistance of stem I in a specific orientation.

Therefore, we hypothesize that TBA initially swings stem I in the solution to catch a freely diffusing tRNA with the help of Specifier, suggesting it follows the ‘fly-casting’ mechanism.^49^ Following this step, the tRNA-bound stem I docks on stem II, which is stabilized by the intra- and inter-backbone hydrogen bonds.

### Stem I Catches tRNA Using the Fly-casting Mechanism

The Specifier from stem I binds to the Anticodon of tRNA at low Mg^2+^ concentrations (≈ 4 mM) through three canonical base pairs. The 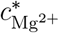 profiles also show that in states such as B1 and B2, where the tRNA is only bound to stem I, the Specifier region does not show significant Mg^2+^ condensation (Fig. 5c) suggesting that, unlike docking, the formation of Specifier-Anticodon hydrogen bonds is relatively less dependent on Mg^2+^. In states B1 and B2, Specifier-Anticodon hydrogen bonds are the only tertiary interactions that stabilize tRNA-T-box binding. Therefore, Specifier-Anticodon hydrogen bond formation is strongly sequence-dependent. To verify the sequence dependence of the tRNA binding affinity, we altered the native Specifier sequence from AUC to CUA by mutating A16C and C18A and performed simulations. We observed that even at [Mg^2+^] = 100 mM, tRNA could not bind to stem I and TBA hopped between states U2 and U3 (Fig. S10), suggesting that the primary Specifier-Anticodon interaction is strongly sequence dependent. Interestingly, despite destabilizing the Specifier-Anticodon interaction, we still did not observe tRNA binding to the S-turn even at [Mg^2+^] = 100 mM. This reinforces the hypothesis that stem I serves as the primary binding site, while the tRNA−S-turn interactions function as an auxiliary but crucial stabilizer for the docking of the tRNA-bound stem I to the S-turn in a specific orientation. This substantiates that the general mechanism of tRNA recognition by TBA is guided by the ‘fly-casting’ principle^49^ (Movie S1).

### tRNA Docking to S-turn Maintains a Specific Orientation

In the native state, tRNA is bound to TBA in such a way that it is close to Ψ.^27^ Simulations show that when tRNA is bound only to stem I (states B1 and B2), the tRNA-bound stem I samples a wide range of conformations due to the scissoring motion of stem I and stem II. To quantify the scissoring motion in tRNA-bound states, we computed the scissoring angle (*ϕ*) between stems I and II (see Data Analysis in SI, Fig. S11). For states B1 and B2, *ϕ* shows broad distributions that support a wide range of motion between stems I and II. The narrow distribution of *ϕ* in the docked state (B3) indicates the restricted motion of stem I. However, *ϕ* cannot detect the structural motion of the docked tRNA. Trajectories showed that after docking to the S-turn, the motion of the tRNA becomes highly restricted to a specific orientation in B3. To quantify the orientation preference of the tRNA with respect to the TBA, we computed an angle, *θ*, between the axis of Ψ−stem II coaxial stacking and the axis of the tRNA Anticodon stem for B3 (Fig. 6). The angle distribution is narrow with an average value ≈ 52^*°*^(± 10^*°*^) independent of [Mg^2+^]. The narrow distribution of *θ* suggests that the tRNA prefers an orientation where it is close to Ψ. This proximity of tRNA to the Ψ enhances the probability of AntiS-tRNA binding (Fig. S12), responsible for checking the aminoacylation status of tRNA. The invariance of this angle under varying [Mg^2+^] indicates that although Mg^2+^ ions contribute to docking stability, the geometry of the tRNA−S-turn inter-backbone hydrogen bonds, and the unique Specifier-Anticodon hydrogen bonds collectively dictate the specific orientation of the docked tRNA.

**Figure 6:**
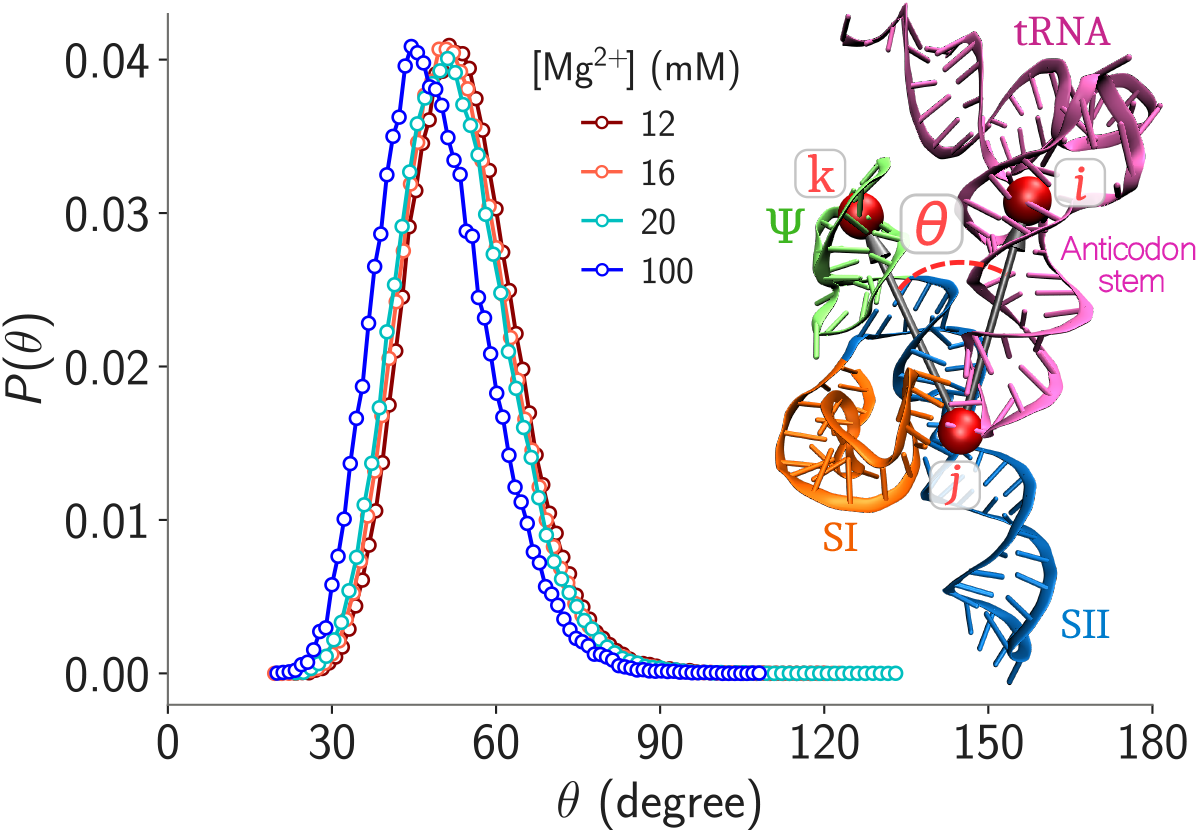
Probability distribution of the orientation angle, *θ*, of the native state (B3) for different [Mg^2+^]. The angle *θ* is calculated using the positions of the beads *i, j*, and *k* shown as red beads. The positions of the beads *i, j*, and *k* are the center of mass of the following beads – **i**: U6, C11, U21, G44 (TBA) **j**: C18, G37, A38 (TBA) and G35, A36 (tRNA) and **k**: U82, A84, G85, C96 (TBA).

## Discussion

Understanding RNA-RNA interactions is critical to deciphering how T-box riboswitches regulate gene expression to regulate the availability of amino acids for protein synthesis in bacteria. In this study, we investigated the folding dynamics of *N. farcinica ileS* T-box riboswitch and its mechanism of tRNA binding using computer simulations with a coarse-grained RNA model.^36^ We validated the RNA model by quantitatively comparing the scattering profiles and radius of gyration of the *apo* T-box riboswitch from simulations and experiments.^23^ Our results complement existing experimental findings and provide new testable predictions for experiments that advance our understanding of tRNA recognition by the T-box riboswitch for gene regulation.

Isothermal titration calorimetry (ITC), SAXS, and smFRET experiments have been performed to probe the conformational dynamics of translational T-box riboswitches across different organisms. ^23,26–28,34,50,51^ The smFRET experiments showed that the T-box riboswitch exhibits a diverse array of slowly interconverting conformations that are coupled to the tRNA binding. smFRET experiments suggested that, in the absence of tRNA, the T-box broadly samples three distinct conformations − open, semi-open, and closed. Our simulations identify the three states (U1, U2, and U3) with detailed structures. We further showed that the varying [Mg^2+^] can redistribute the population of these states. However, without tRNA binding, the docked state is not the global stable state. This finding is also in agreement with the experiments, where a low population of the docked state (≈10%) was observed. Experiments showed that tRNA can directly bind to conformations in the three states of TBA (U1, U2, and U3). Our simulations complement this observation by showing multiple binding and unbinding events (U1 ⇌ B1 and U2 ⇌ B2). However, we did not observe binding of tRNA to TBA conformations in U3. This is attributed to two factors − (i) the low population of U3 in the simulations, and (ii) the inaccessibility of tRNA to the hindered RBD in the U3. smFRET experiments also reported that binding of tRNA to TBA in U3 (U3→B3) is rare. We also found the existence of a new state, B4, which is in equilibrium with the native state (B3).

Mg^2+^ ions strongly influence the folding of TBA and the binding of tRNA.^28^ Simulations showed that Mg^2+^ are critical in stabilizing the following structures: (i) coaxial stacking of Ψ and stem II, restricting the rapid scissoring motion of stems I and II with respect to each other (ii) the hinge at the K-turn region of stem I to facilitate the formation of a predocked state, irrespective of the presence of tRNA and (iii) the tRNA docked structures by neutralizing the electrostatic repulsions between the RNA chains in three different regions (the Specifier from stem I, the S-turn from stem II, and the Anticodon stem loop of tRNA). We have summarized our proposed mechanism of the tRNA recognition process in Fig. 7 by combining all our simulation observations and reported experiments.^23,28,34^

**Figure 7:**
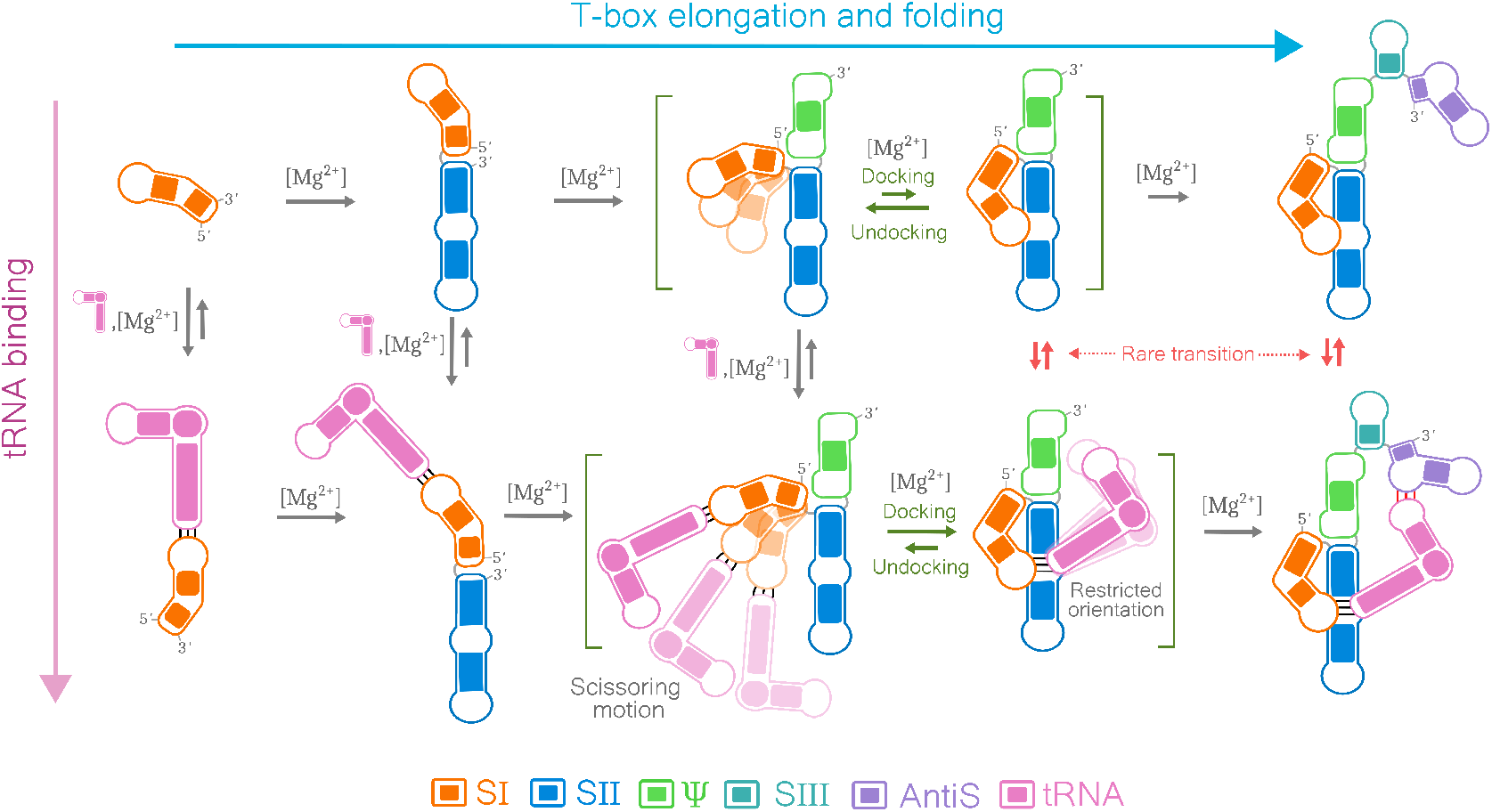
The proposed mechanism of tRNA recognition by *ileS* T-box riboswitch during co-transcriptional folding. The horizontal direction shows the elongation and folding of different domains of the T-box during transcription. The vertical axis shows tRNA binding to different lengths of the folded riboswitch during the transcription process. Specifier-Anticodon interactions are shown with black lines, and tRNA-AntiS interactions are shown with red lines.

RNA starts folding during its transcription, a process known as co-transcriptional folding. The experiments suggest that the tRNA binding takes place in the realm of co-transcriptional folding of the T-box riboswitch (Fig. 7). Our simulations showed that tRNA can bind to the stem I Specifier without any assistance from stem II and Ψ. Since during the transcription stem I gets transcribed first, tRNA can bind to it as soon as it is synthesized. However, after the formation of stem II, stem I prefers to form a coaxial stacking with stem II, which disfavors the native-like docking of stem I and II.^23^ The synthesis of Ψ creates a competition between stem I and Ψ for the coaxial stacking with stem II (Fig. 7). Previous studies have shown that a pseudoknot can also form a stable coaxial stack with helix.^52,53^ The 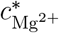 profile showed wherever the Ψ-Stem II coaxial stacking is present (states U2, U3, B2, and B3), there is an enhanced condensation of Mg^2+^ ions around the Ψ-Stem II junction (Fig. 5a-c). This suggests that Mg^2+^ ions stabilize the coaxial stacking of the nascent Ψ pseudoknot and stem II, which facilitates the docking of stem I to stem II.

During the co-transcriptional synthesis of TBA, after the synthesis of Ψ, the discriminator domain is synthesized, which binds to the tRNA and detects its aminoacylation status to regulate the downstream gene. In *N. farcinica ileS* TBA (PDB ID: 6UFM^27^), we observed that the docked tRNA maintains a specific orientation, leaning towards the Ψ. When we compared the crystal structure of TBA with the full structure of *Mycobacterium tuberculosis* (*Mtb*) *ileS* T-box riboswitch aptamer (PDB ID: 6UFH^25^), we found that both form a similar docked state with the tRNA (Fig. S12). In both native structures, tRNA remains oriented in the same direction, regardless of the presence of the discriminator domain. This indicates that docking restrains the orientation of tRNA in such a way that the nascent discriminator domain can immediately interact with the tRNA and make the regulatory decision for the downstream gene. The smFRET experiments on *Mtb ileS* T-box riboswitch showed the existence of two states where the docked tRNA is either bound or unbound to the discriminator domain. In the unbound state, only the discriminator domain changes its orientation and moves away from the tRNA, while the tRNA is stable in its docked orientation.^34^ Similarly, for the structurally different *glyQS* and *tyrS* T-box riboswitches from *B. subtilis*, the tRNA gets anchored at three different regions of the riboswitch.^19,21^ These anchors maintain the directionality of the tRNA toward the discriminator to facilitate discriminator-tRNA binding. The co-crystal structures of the minimal T-box–tRNA complex and cryo-EM studies demonstrate how the curved architecture of the *glyQS* T-box mRNA operates like a mechanical clamp, securing its specific uncharged tRNA to establish the antitermination complex.^19,26^ We conclude that tRNA binding is strongly coupled with the co-transcriptional folding of *ileS* T-box riboswitch, and the orientation restraining of bound-tRNA is an evolutionary strategy of T-box riboswitches to facilitate gene regulation.

Multiple attempts have been made to target the T-box riboswitches for antibiotic development.^29–31,33,54^ Previously, oxazolidinones were used to target the antiterminator domain of the T-box riboswitch to develop new antibacterials. In Gram-positive bacteria, oxazolidinones compete with tRNA to bind to the antiterminator, inhibiting transcription antiter-mination, leading to antibacterial activity.^30,31,54^ In this work, we demonstrated that the initial tRNA interaction with the Specifier region is through the ‘fly-casting’ mechanism, and designing molecules that bind to the specifier region will disrupt this mechanism and the functioning of the riboswitch. Recent reports showed that PKZ18 analogs in combination with aminoglycoside antibiotics targeted the Specifier region of the T-box with high efficacy.^29,33^ We further showed that for *iles* T-box riboswitch, the docking to the S-turn essentially orients the Specifier-bound tRNA towards the discriminator domain to check the aminoacylation state of the tRNA. Therefore, we predict that molecules that can bind to the S-trun and inhibit the docking process have a strong potential to be a drug for *iles* T-box inhibition, irrespective of the tRNA−Specifer binding.

## Conclusion

Taken together, our study reveals that the folding and tRNA-binding dynamics of the *N. farcinica ileS* T-box riboswitches are coupled with co-transcriptional folding, Mg^2+^-mediated stabilization, and competitive coaxial stacking interactions involving stem I, stem II, and the Ψ pseudoknot.^23,28^ This bistable stacking mechanism, which avoids exposing vulnerable single-stranded regions, offers a structurally robust yet conformationally dynamic strategy for RNA regulation.^19,23^ On the other hand, we show that TBA uses the ‘fly-casting’ mechanism to bind tRNA, known for DNA-protein,^49,55^ protein-drug^56^ binding interactions. Thus, combining the structural modularity, bistable stacking, and ‘fly-casting’, the T-box riboswitch serves as a model for understanding how modular RNA elements can orchestrate complex regulatory functions through dynamic yet stable structural ensembles. Additionally, the restrained orientation of docked tRNA across different T-box architectures suggests an evolutionarily conserved mechanism to facilitate rapid engagement with the discriminator domain.^19,26^ The conformational plasticity of these ensembles presents a promising target for therapeutic intervention. Our findings not only deepen the mechanistic understanding of T-box riboswitches but also contribute to the broader framework of RNA-based regulation and drug discovery.

## Supporting information

Movie Showing the Conformational Transitions

## Supporting Information

Simulation details and data analysis; Coarse-grained parameters (Table S1 to S5); Crystal structure of full *M. tuberculosis ileS* (Fig. S1); Trajectory in the absence of tRNA (Fig. S2); Distribution of Inter-arm distances as function of [Mg^2+^] (Fig. S3); Intra-backbone hydrogen bonds (Fig. S4); Trajectories in the presence of tRNA (Fig. S5); tRNA binding domain (RBD) (Fig. S6); FES at different [Mg^2+^] in the presence of tRNA (Fig. S7); Inter-backbone hydrogen bonds (Fig. S8); Trajectory and FES in the absence of inter-backbone hydrogen bonds (Fig. S9); Trajectory of a simulation with mutated Specifier (Fig. S10); Distributions of scissoring angle (Fig. S11); Comparison of simulation structure with existing crystal structures (Fig. S12); Movie of tRNA binding by TBA (Movie S1);

## Acknowledgement

GR acknowledges funding from the Science and Engineering Research Board (SERB) through the grant CRG/2023/002817. DM acknowledges the research fellowship from the Indian Institute of Science, Bangalore. We acknowledge the National Supercomputing Mission (NSM) for providing computing resources of “Param Pravega” at IISc and “Param Brahma” at IISER Pune, supported by the Ministry of Electronics and Information Technology (MeitY) and the Department of Science and Technology (DST), Government of India.

## For Table of Contents Use Only

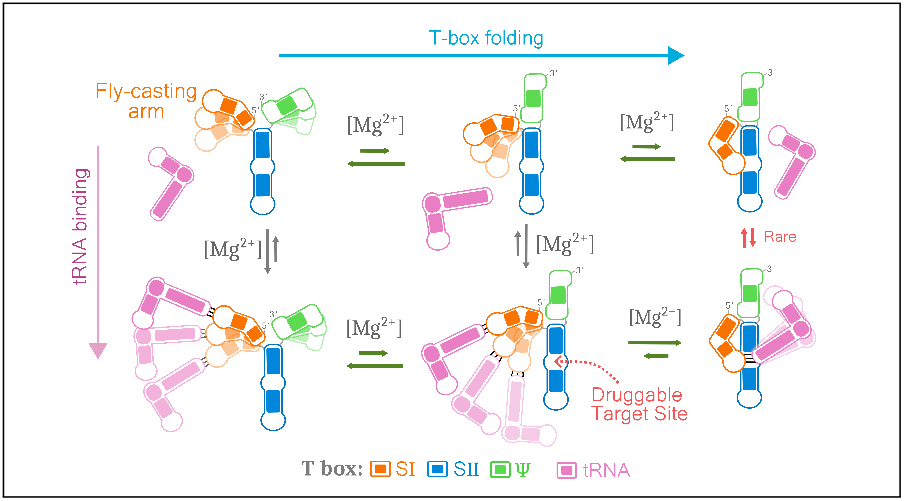

