## Supplementary Information for "Conformational Pathways of Translational T-box Riboswitch Governing tRNA Recognition for Gene Regulation"

### Coarse-grained (CG) simulations

#### Three-Interaction-Site (TIS) Model

We performed the CG simulations using the three interaction site (TIS) RNA model.[1,2] In the TIS model, a nucleotide is modeled using three beads positioned at the center of mass of the phosphate group (P), sugar (S), and nucleobase (B). The TIS energy function includes potentials to model bond length ($U_{\mathrm{BL}}$), bond angle ($U_{\mathrm{BA}}$), excluded volume interactions ($U_{\mathrm{EV}}$), stacking between consecutive bases ($U_{\mathrm{ST}}$), tertiary stacking (stacking between non-consecutive bases) ($U_{\mathrm{TST}}$), electrostatic interactions ($U_{\mathrm{EL}}$) and native hydrogen bond ($U_{\mathrm{HB}}$) interactions present in the folded structure. The energy function[1] is given by

$$U_{\mathrm{TIS}} = U_{\mathrm{BL}} + U_{\mathrm{BA}} + U_{\mathrm{EV}} + U_{\mathrm{EL}} + U_{\mathrm{ST}} + U_{\mathrm{HB}} + U_{\mathrm{TST}} \tag{S1}$$

The bond length and bond angle interactions are modeled using harmonic potentials given by,

$$U_{\mathrm{BL}} = k_\rho(\rho - \rho_0)^2 \tag{S2}$$

$$U_{\mathrm{BA}} = k_\alpha(\alpha - \alpha_0)^2 \tag{S3}$$

where $\rho_0$ and $\alpha_0$ are the bond distance and bond angles at equilibrium from the ideal A-form RNA helix. The $k_\rho$ values are 23, 64, and 10 in kcal.mol$^{-1}$·Å$^{-2}$ for P→S, S→P (' →' implies 5′→3′) and S−B bonds, respectively. The $k_\alpha$ values are 5 kcal mol$^{-1}$ rad$^{-2}$ for angles involving the base beads, and 20 kcal mol$^{-1}$ rad$^{-2}$ otherwise.

The potential used for modeling the RNA hydrogen bonding network is given by,

$$U_{\mathrm{HB}} = U^0_{\mathrm{HB}} \exp(-u) \tag{S4}$$

where,

$$u = 5.0(r-r_0)^2+1.5\{(\theta_1-\theta_{1,0})^2+(\theta_2-\theta_{2,0})^2\}+0.15\{(\psi_0-\psi_{0,0})^2+(\psi_1-\psi_{1,0})^2+(\psi_2-\psi_{2,0})^2\}.$$

The value of $U^0_{\mathrm{HB}}$ is -2.9 kcal·mol$^{-1}$. The parameters $r_0$, $\theta_{1,0}$, $\theta_{2,0}$, $\psi_{0,0}$, $\psi_{1,0}$, and $\psi_{2,0}$ for all the hydrogen bonds are calculated from the coarse-grained model of the riboswitch crystal structure (PDB-ID: 6UFM).[1–3] The canonical hydrogen bonds used in the force field are shown in the secondary structures (see Fig. 1c in the Main text). The other hydrogen bonds and tertiary stacking interactions are in Tables S2, S3, S5, S4. The definitions of $r$, $\theta_1$, $\theta_2$, $\psi_0$, $\psi_1$, and $\psi_2$ are provided in the work of Denesyuk et al.[2] To identify the hydrogen bond network in the crystal structure of the riboswitch, we have used the RNApdbee 2.0 server,[4] and the *N. farcinica ileS* T-box riboswitch aptamer crystal structure (PDB ID: 6UFM).[3]

The stacking interactions between two consecutive nucleotides are sequence-dependent ($U_{\mathrm{ST}}$) and are given by

$$U_{\mathrm{ST}} = \frac{U^0_{\mathrm{ST}}}{1 + k_r(r-r_0)^2 + k_\phi(\phi_1-\phi_{1,0})^2 + k_r(\phi_2-\phi_{2,0})^2} \tag{S5}$$

where, $k_r$ is 1.4 Å$^{-2}$ and $k_\phi$ is 4.0 rad$^{-2}$. The $r_0$, $\phi_{1,0}$, and $\phi_{2,0}$ are calculated from the coarse-grained structure of an ideal A-form RNA helix. The values of $U^0_{\mathrm{ST}}$ and all the definitions of $r$, $\phi_1$, and $\phi_2$ are defined in the original work of Denesyuk et al.[1]

The tertiary stacking interaction between two non-adjacent nucleotide bases ($U_{\mathrm{TST}}$) is given by

$$U_{\mathrm{TST}} = \frac{U^0_{\mathrm{TST}}}{1+u} \tag{S6}$$

where,

$$u = 5.0(r-r_0)^2+1.5\{(\theta_1-\theta_{1,0})^2+(\theta_2-\theta_{2,0})^2\}+0.15\{(\psi_0-\psi_{0,0})^2+(\psi_1-\psi_{1,0})^2+(\psi_2-\psi_{2,0})^2\}.$$

The value of $U^0_{\mathrm{TST}}$ is taken to be -6.5 kcal·mol$^{-1}$. The definitions of the other variables in the potential are the same as the hydrogen bonding interaction potential used in the work of Denesyuk et al.[2]

The electrostatic interaction between the phosphate beads is computed using the Coulomb potential given by.

$$U_{\mathrm{EL}} = \frac{Z_i Z_j}{4\pi\epsilon_0\epsilon_r r_{ij}} \tag{S7}$$

The dielectric constant of water is a function of temperature ($\epsilon_r(T)$) and is given by

$$\epsilon_r(T) = 87.740 - 0.4008T + 9.398 \times 10^{-4}T^2 + 1.410 \times 10^{-6}T^3 \tag{S8}$$

A modified LJ potential is used to model the excluded volume interactions between the beads and is given by

$$U_{\mathrm{EV}} = \begin{cases} \epsilon_{ij}\left[\left(\frac{d_s}{r + d_s - d_{ij}}\right)^{12} - 2\left(\frac{d_s}{r + d_s - d_{ij}}\right)^{6} + 1\right] & \text{if } r \leq d_{ij} \\ 0 & \text{if } r > 0 \end{cases} \tag{S9}$$

where $d_s$ is the diameter of the smallest ion present in the system, $d_{ij} = R_i + R_j$, $\epsilon_{ij} = \sqrt{\epsilon_i \epsilon_j}$. The values of $R_i$ and $\epsilon_i$ for phosphate (P), sugar (S), and bases (A, G, C, and U) are taken from the work of Denesyuk et al.[2] The values of $R_i$ and $\epsilon_i$ for the ions are given in Table S1.

#### CG Simulation Details

We used the three interaction site (TIS-RNA)[1,2] model to study the folding of the *N. farcinica ileS* T-box riboswitch. The initial structure for the simulations was prepared using the crystal structure of the *iles* T-box riboswitch decoder domain (PDB ID: 6UFM).[3] We performed Langevin dynamics simulations using OpenMM[5] at temperature ($T$) 298 K using a cubic box of length 200 Å. The water solvent is implicitly modeled. Simulations were performed at various ion concentrations. In the absence of tRNA, the simulations of TBA were performed in [$Mg^{2+}$] = 2, 4, 6, 8, 10, 20, 40, and 100 mM with a fixed background [$K^+$] = 150 mM. In the presence of tRNA, the simulations of TBA were performed in [$Mg^{2+}$] = 4, 8, 12, 16, 20 mM with a fixed background [$K^+$] = 150 mM. The $K^+$, $Mg^{2+}$, and $Cl^-$ ions were randomly inserted into the box to maintain the required concentration. We performed 8 independent $\approx$ 11 $\mu$s long simulation runs for each concentration. We discarded the initial 1 $\mu$s of the data from each CG trajectory for analysis. The equation of motion for the particles in the simulation is given by

$$m_i \ddot{\vec{r_i}} = -m_i \gamma \dot{\vec{r_i}} + \vec{F_i} + \vec{\Gamma}_i \tag{S10}$$

where, $\vec{r_i}$ and $\gamma$ are coordinates and friction coefficient of the $i$th particle, respectively. The deterministic force on the particle $i$ is given by $\vec{F_i} = -\dfrac{\partial U_{\mathrm{TIS}}(\vec{r_i})}{\partial \vec{r_i}}$, and $\vec{\Gamma}_i$ is an uncorrelated random force with a white noise spectrum. The autocorrelation function of the random force in discretized form is $\left\langle \vec{\Gamma}(t) \cdot \vec{\Gamma}(t+nh) \right\rangle = \frac{2\gamma m_i k_{\mathrm{B}} T}{h} \delta_{0,n}$, where $n = 0, 1, ...$ and $\delta_{0,n}$ is the Kronecker delta function. We have used the *LangevinIntegrator* module in OpenMM to integrate the equations of motion. For better conformational sampling, we used a low friction coefficient ($\gamma = 0.01$ ps$^{-1}$)[6] with an integration time step of 2 fs.

We applied a harmonic potential with a low force constant ($k_s$ = 2 kcal mol$^{-1}$ Å$^{-2}$) between the center of mass (COM) of tRNA and the COM of TBA. This potential is activated only when the distance between the COMs exceeds the cutoff distance ($r_s^{\mathrm{cut}}$ = 85 Å) and remains inactive otherwise. $r_s^{\mathrm{cut}}$ is approximately 3 times the experiential radius of gyration of the TBA ($R_g^{\mathrm{TBA}} \approx$ 29 Å), which ensures that there is enough distance between the TBA and the tRNA in the unbound state and the potential does not bias the binding mechanism. The Particle Mesh Ewald (PME) algorithm was used to compute the long-range Coulomb interactions. We used the TIS2AA program[7] to generate all-atom representations of the coarse-grained structures. We used VMD[8] for visualization and representation of the conformations.

### Data Analysis

#### Fraction of native Contacts

We computed the fraction of native contacts (NC) between a set of nucleotides ($\lambda$) in a conformation $i$ using

$$f_{\mathrm{NC}}^{\lambda,(i)} = \frac{N_{\mathrm{NC}}^{\lambda,(i)}}{N_{\mathrm{NC}}^{\lambda,(\mathrm{cry})}} \tag{S11}$$

where $N_{\mathrm{NC}}^{\lambda,\,\mathrm{cry}}$ and $N_{\mathrm{NC}}^{\lambda,\,(i)}$ are the number of native contacts present between the nucleotides belonging to set $\lambda$ in the crystal structure and the $i^{\mathrm{th}}$ conformation, respectively.[9,10] If the distance between a pair of beads is less than the cutoff distance $r_{cut}$ (= 15 Å),[2,11] then that pair is considered to be in contact.

#### The Scissoring Angle ($\phi$) Between stems I and II

To quantify the relative scissoring motion of stems I and II, we computed the angle $\phi$ between stem I and stem II in states B1, B2, and B3 (Fig. S11). To calculate $\phi$, we used the center of mass positions (i, j, and k). The center of mass positions were calculated using the sugar backbone beads. For i, we used nucleotides U17, C18 from TBA and G35, A36 from tRNA; for j, we used C29, U30, U31, A76, and A77 from TBA; for k, we used C49, G50, C55, and G56 from TBA (Fig. S11).

#### Free Energy Surface (FES) from CG Simulations

We projected the free energy surface (FES) onto $\chi$ to identify the various thermodynamic states populated during TBA folding. We computed the FES $G(\chi)$ using the equation

$$G(f_{\mathrm{NC}}) = -k_{\mathrm{B}}T \ln\big(P(f_{\mathrm{NC}})\big) \tag{S12}$$

where $k_{\mathrm{B}}$ is the Boltzmann constant, and $P(f_{\mathrm{NC}})$ is the probability distribution of $f_{\mathrm{NC}}$ at a given [$Mg^{2+}$].

#### Computation of Contact maps

To compute the average contact map, we have calculated the pair distances between all the RNA CG beads, excluding the beads that are less than three nucleotides apart from each other in the RNA sequence. If a pair distance is less than the cutoff distance $r_{cut}$ (= 15 Å),[2,11] then it is considered a contact between the given pair of beads. The averaging is performed over all the conformations in the sub-ensemble (U1, U2, and U3).

#### Radius of Gyration of Ion Binding Domain

To investigate the change in size of the TBA in different states with varying [$Mg^{2+}$], we computed the average radius of gyration ($\langle R_g^{\mathrm{TBA}} \rangle$) using the equation

$$R_g^{\mathrm{TBA}} = \left( \frac{\sum_{i=1}^{N} m_i \vec{r}_i^{\,2}}{\sum_{i=1}^{N} m_i} \right)^{1/2} \tag{S13}$$

where $m_i$ is the mass of bead $i$ and $\vec{r}_i$ is the distance of bead $i$ from the center of mass of the TBA beads. $m_i$ values are in Table S1.

#### Local Ion Concentration Around RNA

We computed the local ion concentration ($c_j$) to investigate the condensation of positively charged divalent metal ions around phosphate beads using the equation[2]

$$c_j = \frac{1}{N_{\mathrm{A}} V_c} \int_0^{r_c} \mathrm{d}r \; 4\pi r^2 \; \rho_j(r) \tag{S14}$$

where, $\rho_j(r)$ is the number density of a specific type of ion $j$ at a distance $r$ from the specific RNA bead, $V_c$ is the spherical volume with a radius $r_c$, and $N_{\mathrm{A}}$ is the Avogadro number. The cutoff radius is defined as $r_c^{(\mathrm{CG})} = R_{\mathrm{M}^{n+}} + R_{\mathrm{P}} + \Delta r$ ($\approx 5$ Å), where $R_{\mathrm{M}^{n+}}$ is the radius of the the metal ion $\mathrm{M}^{n+}$ (Table S1), $R_{\mathrm{P}}$ (= 2.1 Å) is the CG radius of a phosphate bead[2] and $\Delta r$ ($= 1.6$ Å) is the margin distance, which also leads to $r_c^{(\mathrm{CG})} \approx 5$ Å. Therefore, we set the cutoff radius $r_c = 5$ Å.

Table S1: RNA[2] and ions[12,13] parameters used in the CG simulations.

| Type | $R_i$ (Å) | $m_i$ (amu) | $\epsilon_i$ (kcal mol$^{-1}$) | $Z_i$ |
|---|---|---|---|---|
| P | 2.1 | 62.971 | 0.2 | -1 |
| S | 2.9 | 131.108 | 0.2 | 0 |
| A | 2.8 | 134.119 | 0.2 | 0 |
| G | 3.0 | 150.118 | 0.2 | 0 |
| C | 2.7 | 110.094 | 0.2 | 0 |
| U | 2.7 | 111.079 | 0.2 | 0 |
| $Mg^{2+}$ | 1.353 | 24.305 | 0.009 | +2 |
| $K^+$ | 1.590 | 39.098 | 0.279 | +1 |
| $Cl^-$ | 2.760 | 35.453 | 0.012 | -1 |

Table S2: Non-canonical base-base, base-sugar, sugar-sugar, and phosphate-sugar H-bonds present in the crystal structure of TBA (PDB ID: 6UFM).[3] Canonical base-base H-bonds are shown in the secondary structure.

| Base-Base Non-canonical | Base-Sugar | Sugar-Sugar | Phosphate-Sugar |
|---|---|---|---|
| G7-A26(2), G37-A69(2), A38-U68(2), A39-A66(2), G41-U64(2), G4-A26(1), A8-G25(1), A8-G27(1), U9-G24(1), G13-C15(1), U17-G70(1), A35-A72(1) C40-C65(1), G67-U68(1), G79-A92(1), G81-C86(1), U82-A84 (1), U82-C96(1), C87-A94(1), G89-U90(1) | A5-A26, A8-G27, A26-G4, A25-G7, G37-G67, A39-A19, C40-C65, A69-C18, A69-C37, G70-U17, C88-A92, A92-G79, C93-C87 | U9-C28, A16-A71, U17-A70, C18-A69, A19-A39, U17-U71 | U17-C15, U30-U90, C93-G80 |

Table S3: Non-canonical base-base, base-sugar, and phosphate-sugar H-bonds present in the crystal structure of tRNA (PDB ID: 6UFM).[3] Canonical base-base H-bonds are shown in the secondary structure.

| Base-Base Non-cannonical (Counts) | Base-Sugar | Phosphate-Sugar |
|---|---|---|
| A9-A24 (2), G18-U56 (1), G23-G47 (2), G27-A45 (2), C33-A39 (1), G46-G10 (1) | A22-U8, G58-G18, G58-G19, A36-U34, G58-U56 | G50-A7, U61-A59 |

Table S4: Canonical base-base, non-canonical base-base, base-sugar, sugar-sugar, and phosphate-sugar hydrogen bonds and the base-base tertiary stacks present between tRNA and TBA in the crystal structure (PDB: 6UFM).[3]

| Base-Base Canonical H-Bonds (Counts) | Base-Base Non-Canonical H-Bonds (Counts) | Base-Sugar | Sugar-Sugar | Phosphate-Sugar |
|---|---|---|---|---|
| $G35^{(tRNA)}$ - $C18^{(TBA)}$ (3), $A36^{(tRNA)}$ - $U17^{(TBA)}$ (2), $U37^{(tRNA)}$ - $A16^{(TBA)}$ (2), | $G35^{(tRNA)}$ - $A38^{(TBA)}$ (1) | $G35^{(tRNA)}$ - $A38^{(TBA)}$, $U37^{(tRNA)}$ - $A71^{(TBA)}$ | $G35^{(tRNA)}$ - $A38^{(TBA)}$, $A36^{(tRNA)}$ - $G37^{(TBA)}$, $A36^{(tRNA)}$ - $A38^{(TBA)}$ | $A39^{(tRNA)}$ - $A72^{(TBA)}$ |

Table S5: Base-base tertiary stacks present in the crystal structure of TBA and tRNA (PDB ID: 6UFM).[3]

| TBA | tRNA |
|---|---|
| G4-G27, G7-G25, A8-A26, G12-G22, G33-G75, A38-A69, G44-G62, G47-G59, G50-G56, G79-G89, G85-U97, A5-A26, G37-G67, A66-U68, U30-G89, U90-A92 | G15-G60, G18-G58, G19-G58, A22-G47, A9-G46, G18-A59, A59-G62 |

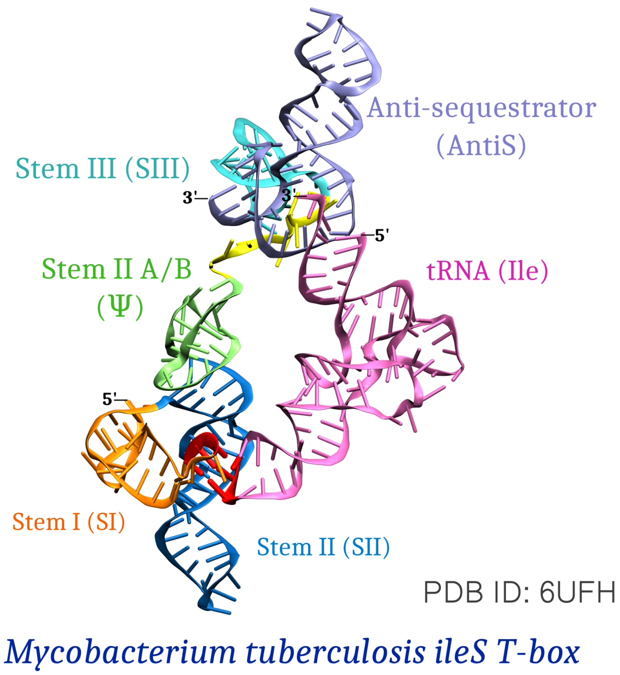


Figure S1: Crystal structure of *M. tuberculosis ileS* T-box riboswitch showing the decoder and discriminator domains (PDB ID: 6UFH)[14]

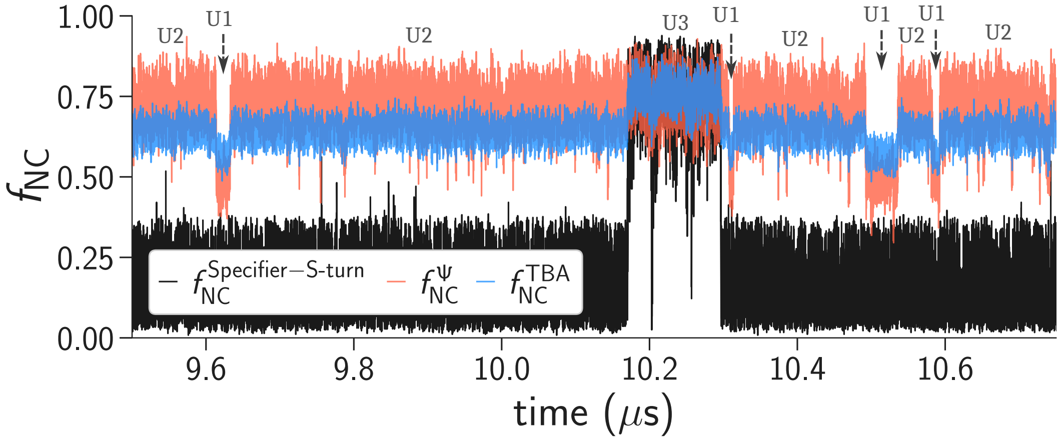


Figure S2: Fraction of native contacts as a function of time in a simulation trajectory showing the transitions between states U1, U2, and U3 in the absence of tRNA.

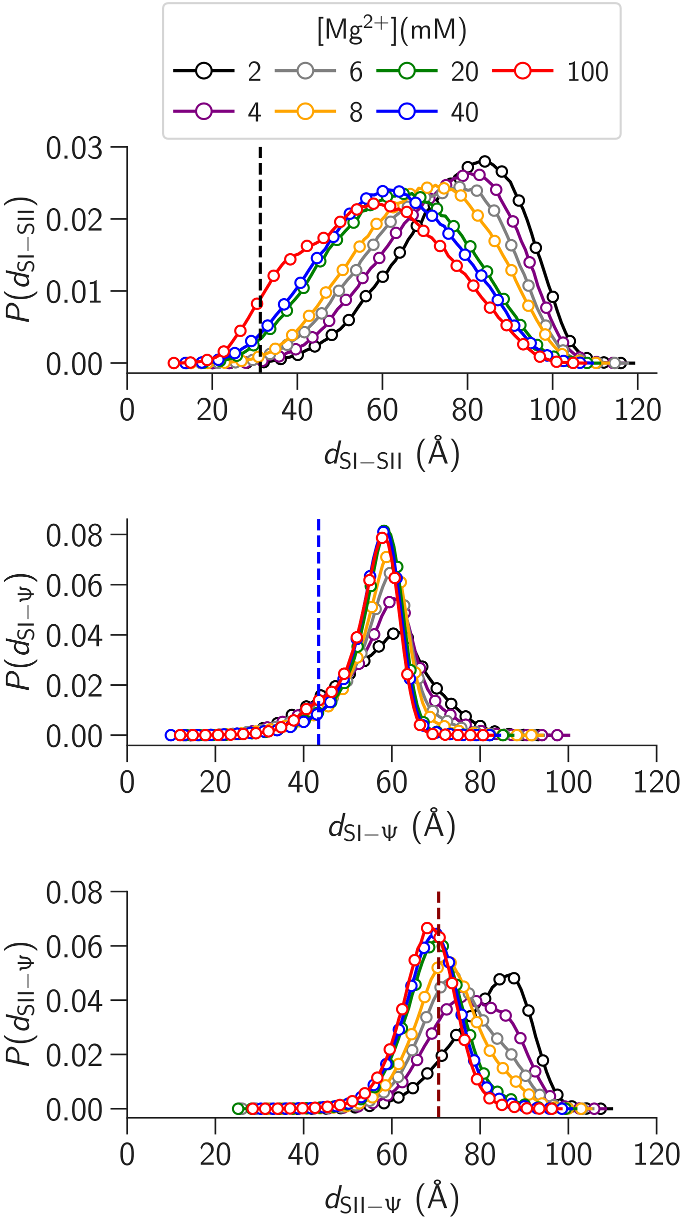


Figure S3: Probability distributions of distances between (a) stem I and stem II, (b) stem I and Ψ, and (c) stem II and Ψ for different [$Mg^{2+}$]. The corresponding distances in the crystal structure (PDB ID: 6UFM)[3] are shown as vertical dashed lines.

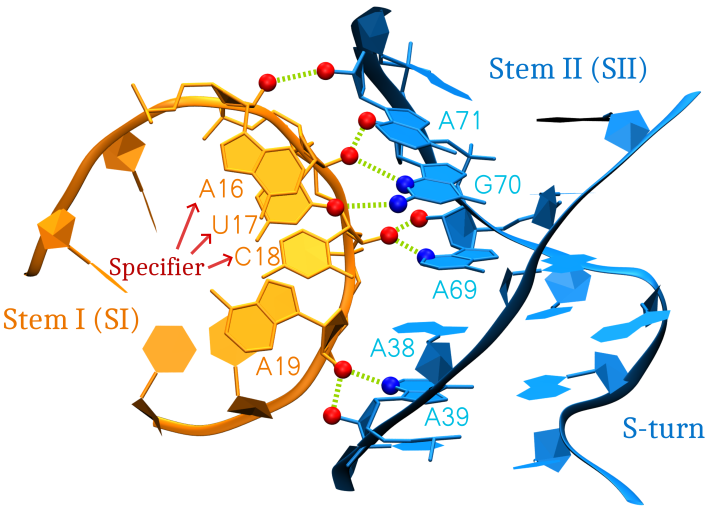


Figure S4: The intra-backbone hydrogen bonds known as the ribose zipper[15] interactions between stems I and II in TBA from the crystal structure (PDB ID: 6UFM) shown in Table S2.[3]

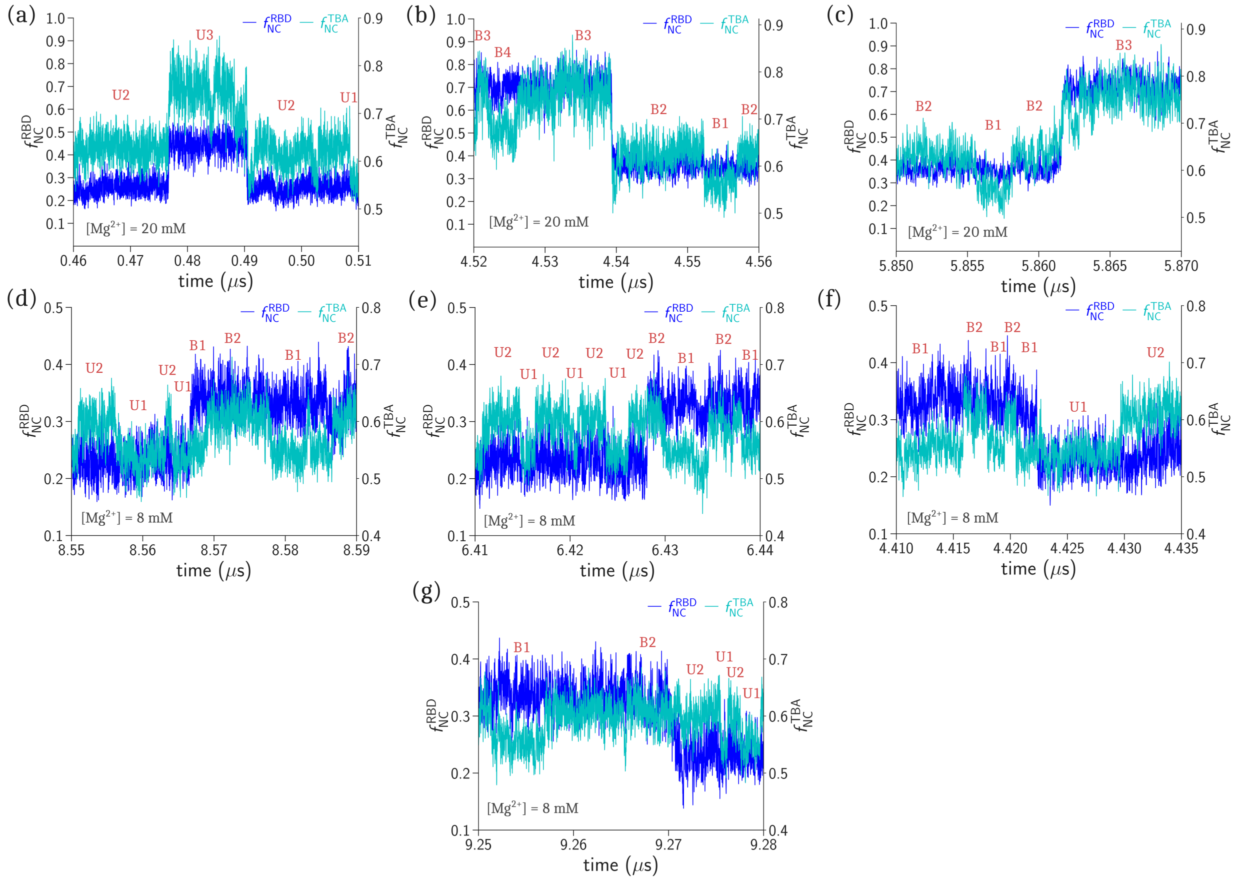


Figure S5: Fraction of native contacts in the TBA, $f_{\mathrm{NC}}^{\mathrm{TBA}}$ (right axis) and the fraction of native contacts in the RBD, $f_{\mathrm{NC}}^{\mathrm{RBD}}$ (left axis) (Fig S6) as function of time in representative trajectories showing transitions between the states U1, U2, U3, B1, B2, B3, and B4.

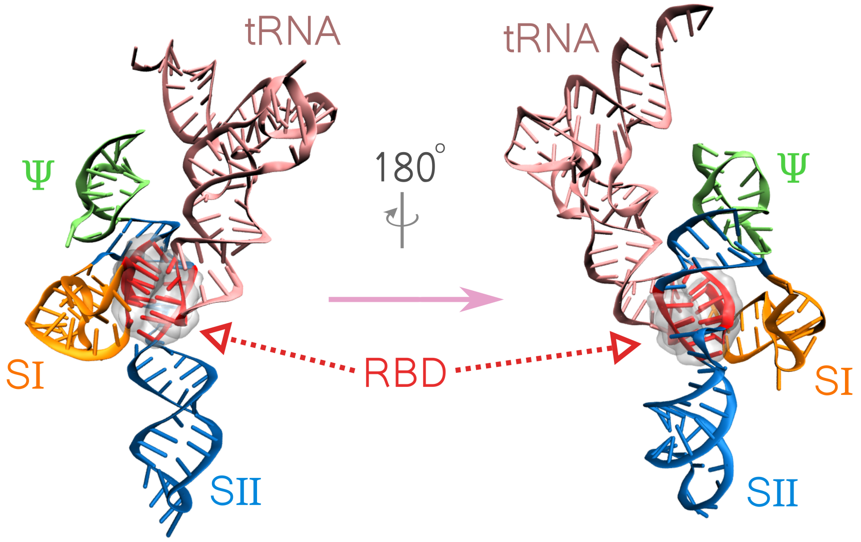


Figure S6: Nucleotides A16-A19, A36-A39, A69-A72 from TBA, and G35-A38 from tRNA constitute the tRNA binding domain (RBD) and are shown in red.

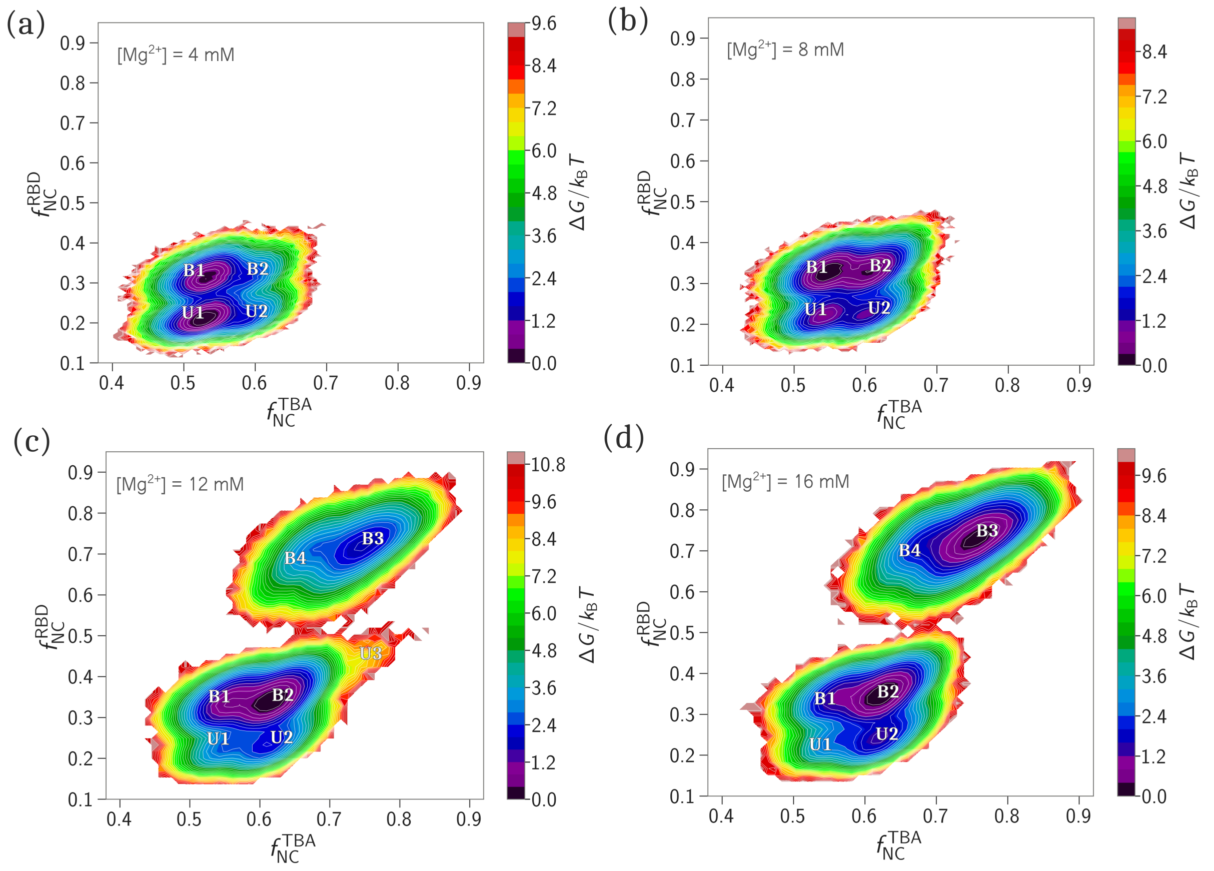


Figure S7: Two dimensional FES projected onto $f_{\rm NC}^{\rm TBA}$ and $f_{\rm NC}^{\rm RBD}$ shows distinct states for $[{\rm Mg}^{2+}]$ = **(a)** 4 mM, **(b)** 8 mM, **(c)** 12 mM, and **(d)** 16 mM.

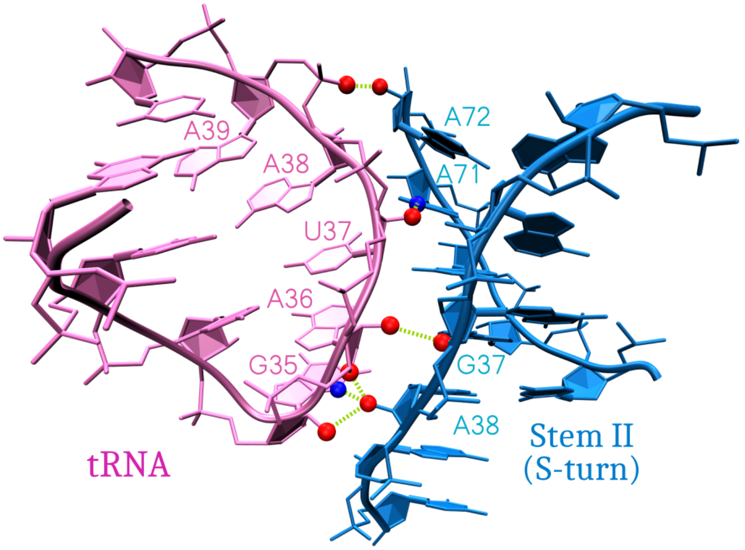


Figure S8: The inter-backbone hydrogen bonds between tRNA and TBA from the crystal structure (PDB ID: 6UFM) shown in Table S4.[3]

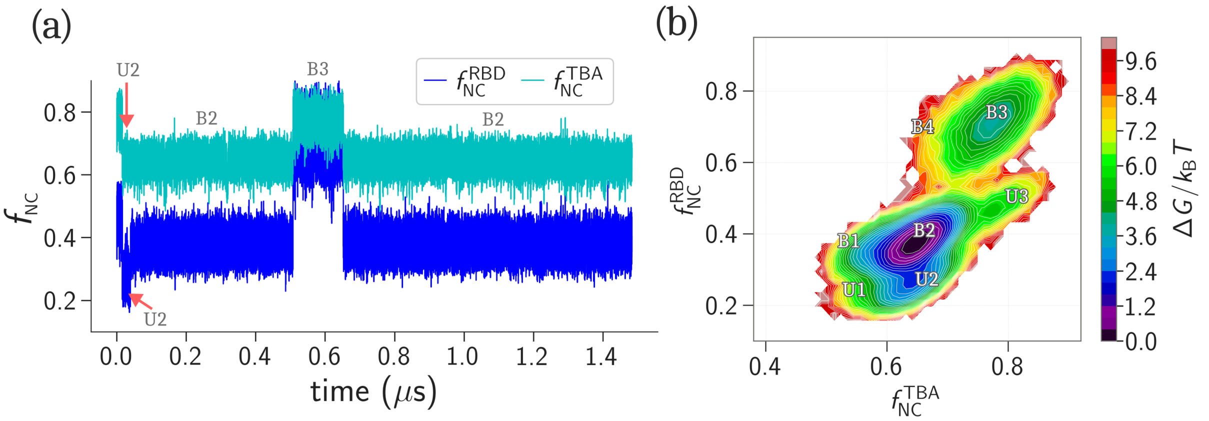


Figure S9: (a) Fraction of native contacts in TBA ($f_{NC}^{TBA}$) and fraction of native contacts in RBD ($f_{NC}^{RBD}$) (Fig S6) as a function of time in a representative trajectory with no inter-backbone H-bonds between tRNA and S-turn for $[Mg^{2+}] = 100$ mM. (b) The 2D free energy surface projected onto $f_{NC}^{TBA}$ and $f_{NC}^{TBA}$.

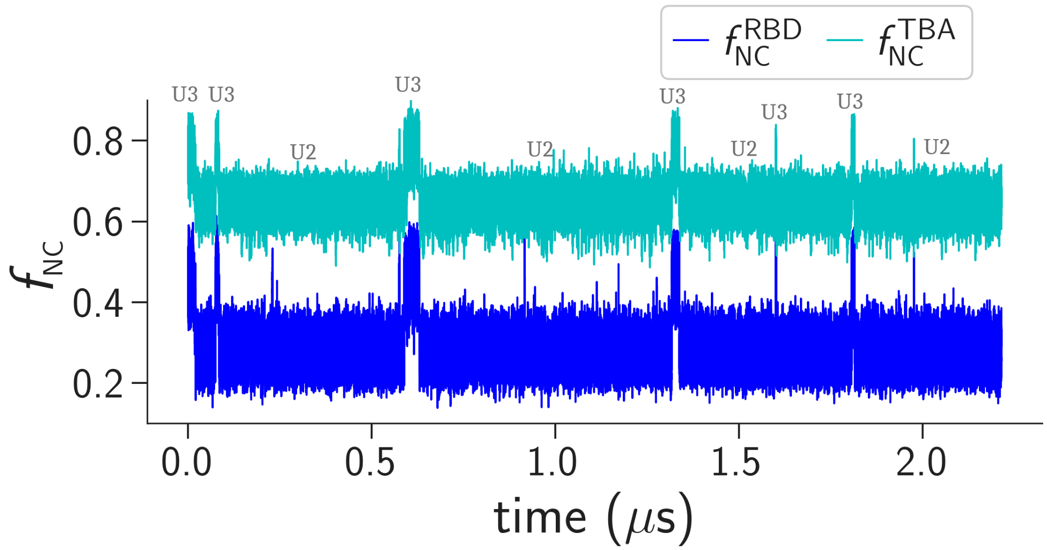


Figure S10: A representative trajectory from the simulations where we altered the Specifier sequence from AUC to CUA by mutating A16C and C18A. The trajectory show fraction of native contacts in TBA ($f_{NC}^{TBA}$) and fraction of native contacts in RBD ($f_{NC}^{RBD}$) (Fig S6) as function of time for [$Mg^{2+}$] = 100 mM. We did not observe any tRNA-stem I binding.

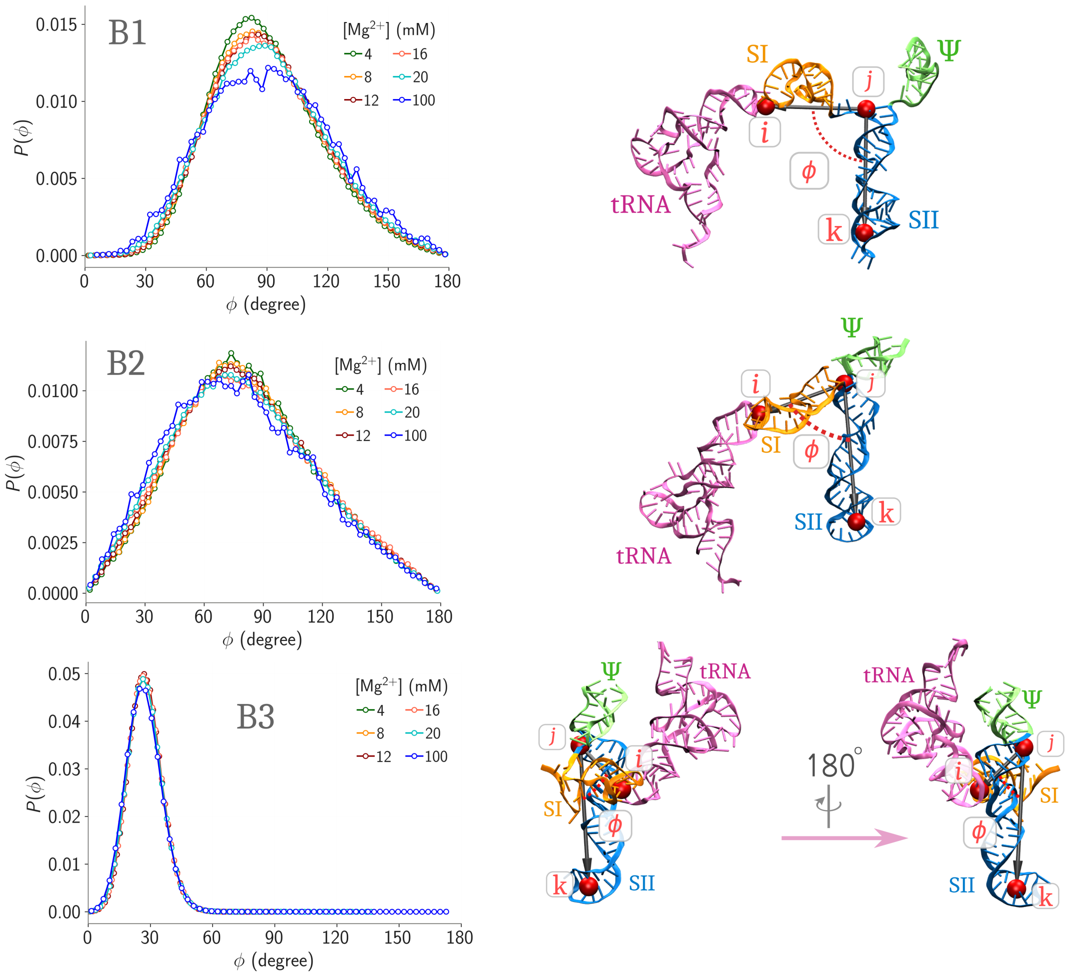


Figure S11: Probability distribution of scissoring angle, $\phi$, in states B1, B2, and B3 for different [$Mg^{2+}$] (left column). The scissoring angle, $\phi$, is shown in representative structures (right column).

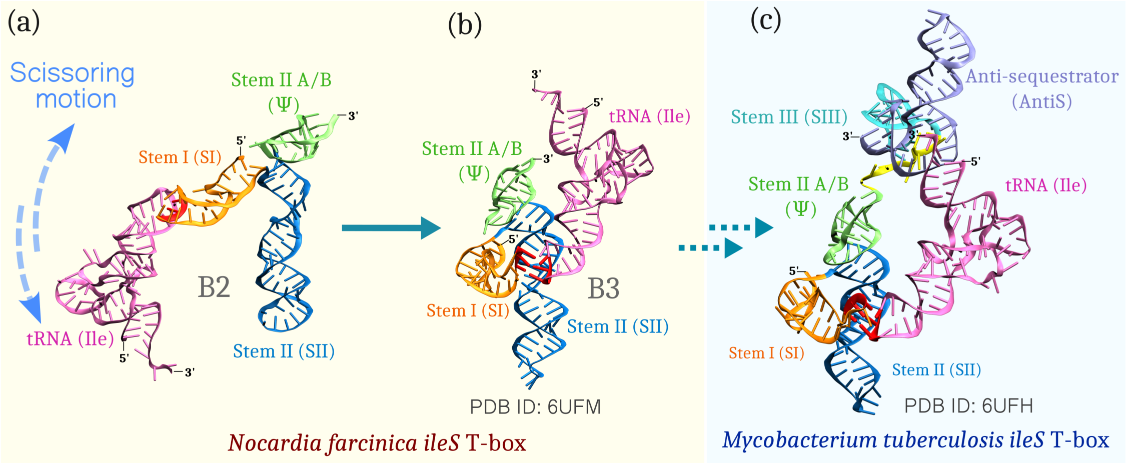


Figure S12: Structural transition in *N. farcinica ileS* T-box riboswitch decoder domain (TBA) from (a) state B2 to (b) state B3 (native state) is shown in the pale-yellow shaded region. (c) The crystal structure of M. tuberculosis ileS T-box riboswitch with both the decoder (SI, SII, and Ψ) and the discriminator (SIII, and AntiS) is shown in the sky-blue shaded region. The decoder and discriminator are joined via a connector sequence (yellow). Comparison of structures (b) and (c) shows that the bound tRNA is in proximity to Ψ, which further facilitates the AntiS-tRNA binding.

#### Movie S1

Binding of a freely diffusing tRNA by a T-box decoder domain (TBA) via a fly-casting mechanism.[16]
